# The interactome of the UapA transporter reveals putative new players in anterograde membrane cargo trafficking

**DOI:** 10.1101/2023.07.21.550021

**Authors:** Xenia Georgiou, Sofia Dimou, George Diallinas, Martina Samiotaki

**Affiliations:** Department of Biology, National and Kapodistrian University of Athens, Panepistimioupolis, Athens, 15784, Greece; Institute of Molecular Biology and Biotechnology, Foundation for Research and Technology, Heraklion, 70013, Greece; Biomedical Sciences Research Center “Alexander Fleming”, Institute for Bioinnovation, Vari, 16672, Greece

**Keywords:** ER-exit, COPII, COPI, traffic, fungi, Golgi-bypass, UapA, Proximity Dependent Biotinylation, proteomics

## Abstract

Neosynthesized plasma membrane (PM) proteins co-translationally translocate to the ER, concentrate at regions called ER-exit sites (ERes) and pack into COPII secretory vesicles which are sorted to the early-Golgi through membrane fusion. Following Golgi maturation, membrane cargoes reach the late-Golgi, from where they exit in clathrin-coated vesicles destined to the PM, directly or through endosomes. Post-Golgi membrane cargo trafficking also involves the cytoskeleton and the exocyst. The Golgi-dependent secretory pathway is thought to be responsible for the trafficking of all major membrane proteins. However, our recent findings in *Aspergillus nidulans* showed that several plasma membrane cargoes, such as transporters and receptors, follow a sorting route that seems to bypass Golgi functioning. To gain insight on membrane trafficking and specifically Golgi-bypass, here we used proximity dependent biotinylation (PDB) coupled with data-independent acquisition mass spectrometry (DIA-MS) for identifying transient interactors of the UapA transporter. Our assays, which included proteomes of wild-type and mutant strains affecting ER-exit or endocytosis, identified both expected and novel interactions that might be physiologically relevant to UapA trafficking. Among those, we validated, using reverse genetics and fluorescence microscopy, that COPI coatomer is essential for ER-exit and anterograde trafficking of UapA and other membrane cargoes. We also showed that ArfA^Arf1^ GTPase activating protein (GAP) Glo3 contributes to UapA trafficking at increased temperature. This is the first report addressing the identification of transient interactions during membrane cargo biogenesis using PDB and proteomics coupled with fungal genetics. Our work provides a basis for dissecting dynamic membrane cargo trafficking via PDB assays.

**Highlights:**

- A PDB-proteomics system for identifying transient interactions in *A.nidulans*
- UapA interactomes in wild-type and specific mutants
- Analysis and validation of UapA interacting proteins
- COPI and Arf1 are essential for cargo anterograde trafficking

**Graphical Abstract:** 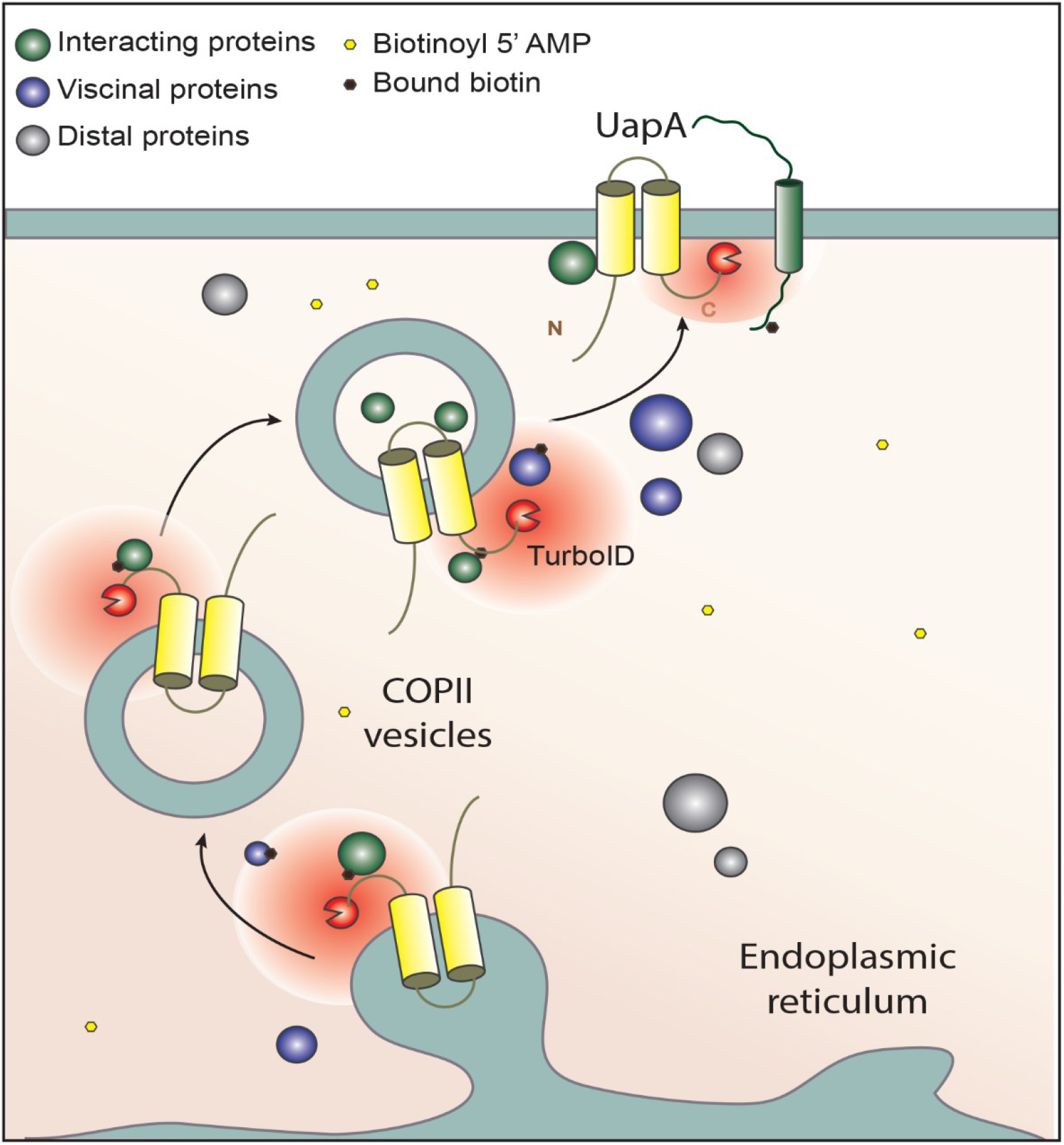

## 1. Introduction

One of the fundamental differences between eukaryotic and prokaryotic cells is their level of compartmentalization. In eukaryotic cells, molecules must be located in specific subcellular compartments to function properly. This vital process is tightly regulated and relies on proper protein folding and distinct physiological signals. Membrane proteins follow the highly conserved membrane-trafficking system to reach their appropriate destinations (Bonifacino and Glick, 2004). Conventionally, membrane proteins are co-translationally translocated to the endoplasmic reticulum (ER), where they are sorted into nascent ER-exit sites (ERes) and packaged into coated COPII secretory vesicles. These vesicles bud and fuse with the early-Golgi (*cis*-Golgi), before reaching the late-Golgi (*trans*-Golgi network or TGN) via Golgi maturation (Emr et al., 2009; Pantazopoulou & Glick, 2019). From the TGN, membrane proteins destined for the plasma membrane (PM) are secreted via AP1/clathrin-coated vesicles, a process regulated by Rab GTPases and requiring microtubule and actin polymerization (Pantazopoulou et al., 2014; Takeshita et al., 2014; Robinson, 2015). Fusion between vesicular intermediates and target membranes is mediated by N-ethylmaleimide-sensitive factor attachment protein receptors (SNAREs), tethering proteins, small GTPases and other regulatory factors (Rothman, 1994; Jahn & Südhof, 1999; Rojas et al., 2012; Gillingham et al., 2014; Wu & Guo, 2015; Pinar & Peñalva, 2021).

Our recent studies in the model fungus *Aspergillus nidulans* have however identified a novel Golgi-independent pathway for trafficking and translocation to the PM of major membrane proteins, such as nutrient transporters, the proton pump ATPase PmaA or the PalI alkaline pH-sensor component (Dimou et al., 2020; Dimou et al., 2022). Interestingly, cargoes bypassing the Golgi still necessitate functional core COPII components (e.g., Sec24, Sec13). This in turn strongly suggests that there must be two subpopulations of COPII vesicles exiting the ER. The first, the conventional one, is sorted to the early Golgi. The second, the one that deviates from the Golgi, is sorted to the PM either directly or via an unknown endosomal compartment. Notably, the Golgi-bypassing cargoes show a clear anti-polar and rather homogenous distribution all over the membrane of *A. nidulans* hyphal cells, whereas conventional cargoes, such as chitin synthase (ChsB), lipid flippases (DnfA and DfnB) or the v-SNARE SynA, are highly polarized cargoes, located in the apex of growing hyphae. What defines the composition of the two different types of COPII vesicles and what route is followed after ER-exit for the vesicles bypassing the Golgi remains unknown. Apparently, the cargoes themselves seem to dictate their own trafficking routes and underlying mechanisms involved. Addressing how functionally distinct membrane cargoes contain the information to follow distinct trafficking routes and dissect the mechanistic details underlying Golgi-bypass will help unveil a critical aspect of a basic cellular process in eukaryotes.

A highly efficient and popular method to identify transient protein interactions *in vivo* is proximity dependent biotinylation (PDB) coupled to mass spectrometry (LC-MS/MS) and targeted proteomics analysis (Branon et al., 2018; Gingras et al., 2019). PDB assays involve a biotin ligase fused in-frame with a protein of interest. The ligase utilizes biotin to catalyze the formation of biotinoyl-5′-AMP, which generates activated biotin molecules capable of reacting with free primary amines (typically lysine residues) of neighboring proteins. This process enables biotinylation of proximal proteins in the native cellular environment. Subsequently, biotinylated proteins are specifically enriched through biotin affinity capture from denatured cell lysates and identified and quantified using liquid chromatography-mass spectrometry (LC-MS/MS). Unlike many other affinity capture approaches for studying protein-protein interactions, PDB does not rely on physical protein-protein binding within native cell lysates. This unique feature allows for the identification of protein proximities that are weak, transient, and dynamic, such as the interactions occurring during vesicular secretion. PDB approaches have been established in animal cells, plants, and recently fungi (for examples see Mair et al., 2019; Larochelle et al., 2019; Arora et al., 2020; Kershberg et al, 2022, Yang et al., 2022; Hollstein et al., 2022; Fenech et al., 2023), but have not been used in *A. nidulans*. Also, to our knowledge, no study has used PDB methods for identifying the very rapid and transient interactions of a moving transmembrane cargo.

Here, we used PDB coupled with Liquid Chromatography with Data-independent acquisition mass spectrometry (DIA-MS) as an unbiased targeted approach for identifying transient protein interactions involved in the trafficking of the UapA xanthine-uric acid transporter (Alguel et al., 2016; Diallinas 2016). UapA was chosen as the most extensively cargo proposed to bypass the Golgi (Dimou et al., 2020; Dimou et al., 2022). To achieve our goals we developed a functional system that makes use of TurboID biotin ligase (Branon et al., 2018) fused to UapA and performed PDB assays with the proteomes of a wild-type (wt) strain and mutants affected ER-exit or endocytosis of UapA. We identified both expected and novel putative interactions that might be physiologically relevant to UapA trafficking. Among the best hits, and taking into account the interactomes of wt and mutant strains, we prioritized to validate the role of CopA, a component of COPI coatomer, Arf^Arf1^ small GTPase, which drives COPI-mediated vesicle budding needed for cargo trafficking, and two ArfA^Arf1^ GTPase activating proteins (GAPs), namely Glo3 and Gcs1.

## Materials and Methods

### 2.1 Media, strains and growth conditions

Standard complete and minimal media for *A. nidulans* were used (FGSC, http://www.fgsc.net). Media and chemical reagents were obtained from Sigma-Aldrich (Life Science Chemilab SA, Hellas) or AppliChem (Bioline Scientific SA, Hellas). Glucose 1% (w/v) or fructose 0.1% (w/v) was used as carbon sources. Nitrogen sources were used at the following final concentrations: sodium nitrate (NaNO_3_) 10 mM, ammonium tartrate (NH_4_^+^) 10 mM, uric acid 0.5 mM, hypoxanthine 0.5 mM and allopurinol 3 μM. Thiamine hydrochloride was used at a final concentration of 10-20 μM as a repressor of the *thiA* promoter (Apostolaki et al., 2012). *A. nidulans* transformation was performed by generating protoplasts from germinating conidiospores as described previously (Koukaki et al., 2003). A Δ*uapA* Δ*uapC::AFpyrG* Δ*azgA pabaA1 argB2* mutant strain, named Δ3 (Pantazopoulou et al., 2007), was the recipient strain in transformations with plasmids carrying TurboID tagged wild-type or mutant versions of UapA. Selection was based on complementation of the arginine auxotrophy *argB2*. An *nkuA* DNA helicase deficient strain (TNO2A7; Nayak et al., 2006) was used as the recipient strain for generating “in locus” integrations of gene fusions with fluorescent tags, promoter replacement or deletion cassettes by the *A. fumigatus* markers orotidine-50-phosphate-decarboxylase (AF*pyrG*, Afu2g0836), GTP-cyclohydrolase II (AF*riboB*, Afu1g13300) or a pyridoxine biosynthesis gene (AF*pyroA*, Afu5g08090), resulting in complementation of the relevant auxotrophies. Transformants were verified by PCR, Western blot and growth test analysis. Combinations of mutations and fluorescent epitope-tagged strains were generated by standard genetic crossing and progeny analysis. Growth tests were performed at 25, 30 or 37°C, at pH 6.8. *A. nidulans* strains used are listed in **Appendix Table S1**.

### 2.2 Nucleic acid manipulations and genetic constructions

Genomic DNA extraction from *A. nidulans* was performed as described in FGSC (http://www.fgsc.net). Plasmids, prepared in *E. coli*, and DNA restriction or PCR fragments were purified from agarose 0.8 % gels with the Nucleospin Plasmid Kit or Nucleospin ExtractII kit, according to the manufacturer‟s instructions (Macherey-Nagel, Lab Supplies Scientific SA, Hellas). Standard PCR reactions were performed using KAPATaq DNA polymerase (Kapa Biosystems). PCR products used for cloning, sequencing and re-introduction by transformation in *A. nidulans* were amplified by a high fidelity KAPA HiFi HotStart Ready Mix (Kapa Biosystems) polymerase. DNA sequences were determined by VBC-Genomics (Vienna, Austria). Vectors expressing UapA-TurboID, UapA-DYDY-TurboID were constructed using as a template the plasmid pAN510, carrying the *uapA* promoter and the *argB* selection marker (Diallinas et al., 1998). Gene cassettes were generated by sequential cloning of the relevant fragments in the pGEM-T plasmid, which served as template to PCR-amplify the relevant linear cassettes for *glo3* and *gcs1* knock-outs, *copA* and *arfA* knock-downs and fluorescent epitope-tagged strains. Forward and reverse primers used are listed in **Appendix Table S2**.

### 2.3. Growth Conditions and Proximity Dependent Biotinylation Assays

Cultures of *A. nidulans* were grown in minimal medium (MM) supplemented with 1% glucose (w/v) and 10 mM NH_4_^+^ (di-ammonium tartrate) at 37°C for 16 h at 140 rpm. To enable the expression of UapA-TurboID or UapA-DYDY-TurboID, the culture was shifted to a minimal medium with 1% glucose, 10 mM NaNO_3_, 50 μM biotin at 30°C for 4 h at 140 rpm. The collected mycelia were washed thoroughly with cold sterilized water to remove the residual biotin and subsequently were snap frozen in liquid nitrogen and pulverized in a mortar with a pestle. For each strain (Δ3, UapA-TurboID, UapA-DYDY-TurboID, Δ*artA*: UapA-TurboID) three biological replicates were prepared, using approximately 700 mg of fine powder from *A. nidulans* mycelium. The powder was dissolved in 3 mL lysis buffer [4% (w/v) SDS, 0.1 M Tris-HCL, 0.1 M dithioerythritol (DTE) pH = 7.6] (Lysis buffer), and lysis took place in a water bath sonication (4 cycles, 30 min each at 45° C) apparatus [Dimou et al., 2021]. After complete solubilization, centrifugation took place for 10 min, at 12,000 rpm and the remaining debris was discarded. The protein extracts collected were stored at -80°C until further processing.

Protein extracts were transferred into an Amicon Ultra Centrifugal filter device (10 kDa MWCO, Merck) for buffer exchange [50 mM Tris-HCL pH 7.5, 150 mM NaCl, 0.1% (w/v) SDS, 1% Triton-X-100, 0.5% Na-deoxycholate, 1 mM EDTA, 1 mM DTE] and protein concentration. This step was crucial for depletion of free biotin from the samples (Mair et al., 2019). The protein concentration of each extract was then measured by Bradford Assay. For each replicate, a volume corresponding to 5 mg protein was transferred into an Eppendorf tube containing Dynabeads MyOne Streptavidin C1 (Invitrogen) from 100 μl bead slurry that were pre-washed twice with extraction buffer (50 mM Tris-HCL pH 7.5, 150 mM NaCl, 0.1% (w/v) SDS, 1% Triton-X-100, 0.5% Na-deoxycholate, 1 mM EDTA, 1 mM DTE). The incubation took place on a rotor wheel at 4°C overnight (∼16 h). The next day, the beads were separated from the protein extract on a magnetic rack and washed with the following solutions: 2x 1 mL cold extraction buffer, 1x 1 mL cold 1 M KCL, 1x cold 100 mM Na_2_CO_3_, 1x 1mL 2M Urea in 10 mM Tris pH 8 at room temperature (Branon et al., 2018).

### 2.4 On-Bead Tryptic Digestion-MS sample preparation

After the discard of the final wash solution, 100 μL of Tryptic digestion Buffer (1 M Urea, 1 mM DTE, 50 mM Tris pH 8.2, 1 μg Trypsin/Lys-C Protease Mix, Thermo) were added and the incubation took place on a thermoshaker at 37°C, 1300 rpm, for 3 h. Then, the supernatant was collected and the beads were washed twice with 25 μL of 1 M urea, 1 mM DTE, 50 mM Tris-HCL pH 8.2. Supernatants were pooled on new tubes and DTE was added to a final concentration of 4 mM on each sample for 30 min, on a thermoshaker at 37°C, 1300 rpm. Iodoacetamide was added to a final concentration of 10 mM on each sample and incubation took place in the dark for 45 min. Additional 0,5 μg of trypsin was added for overnight digestion (12-13 h), at 37°C (Zhang et al., 2019). The next day, the peptides were purified with C-18 desalting 100 µL zip-tips (Pierce™ C18 Tips) according to manufacturer instructions and finally solubilized in the mobile phase A (0.1% Formic acid in water), sonicated and the peptide concentration was determined through absorbance at 280 nm measurement using a nanodrop instrument.

### 2.5 LC-MS/MS

Samples were analyzed on a liquid chromatography tandem mass spectrometry (LC-MS/MS) setup consisting of a Dionex Ultimate 3000 nanoRSLC coupled in line with a Thermo Q Exactive HF-X Orbitrap mass spectrometer. Peptidic samples were directly injected and separated on an 25 cm-long analytical C18 column (PepSep, 1.9μm3 beads, 75 µm ID) using an one-hour long run, starting with a gradient of 7% Buffer B (0.1% Formic acid in 80% Acetonitrile) to 35% for 40 min and followed by an increase to 45% in 5 min and a second increase to 99% in 0.5 min and then kept constant for equilibration for 14.5min. A full MS was acquired in profile mode using a Q Exactive HF-X Hybrid Quadrupole-Orbitrap mass spectrometer, operating in the scan range of 375-1400 m/z using 120K resolving power with an AGC of 3x 106 and maximum IT of 60 ms followed by data independent acquisition method using 8 windows (a total of 39 loop counts) each with 15K resolving power with an AGC of 3x 105 and max IT of 22ms and normalized collision energy (NCE) of 26. Each sample (e.g. three biological replicas) was analyzed in three technical replicas. Orbitrap raw data were analyzed in DIA-NN 1.8.1 (Data-Independent Acquisition by Neural Networks) through searching against the *Aspergillus (Emericella) nidulans* Reference Proteome (downloaded from Uniprot (UP000000560), 10,561proteins entries, downloaded in February 2022) using the library free mode of the software, allowing up to two tryptic missed cleavages and a maximum of three variable modifications/peptide. A spectral library was created from the DIA runs and used to re-analyze them (double search mode). DIA-NN search was used with oxidation of methionine residues and acetylation of the protein N-termini set as variable modifications and carbamidomethylation of cysteine residues as fixed modification. The match between runs feature was used for all analyses and the output (precursor) was filtered at 0.01 FDR and finally the protein inference was performed on the level of genes using only proteotypic peptides. The resulting spectra from each strain (Δ3, UapA-TurboID, UapA-DYDY-TurboID, Δ*artA*:UapA-TurboID) were analyzed with MaxQuant (version 2.1.4.0, www.maxquant.org; Cox and Mann, 2008; Cox et al., 2011). Label free analysis was performed with Perseus Values were log (2) transformed, a threshold of 70% of valid values in at least one group was applied and the missing values were replaced from normal distribution (Tyanova, S. et al. 2016). For statistical analysis, Student‟s t-test was performed, and permutation-based FDR (0.05) was calculated. The mass spectrometry proteomics data have been deposited to the ProteomeXchange Consortium via the PRIDE partner repository with the dataset identifiers: PXD043934, PXD043941, PXD043944 (Perez-Riverol et al., 2022).

### 2.6 MS data analysis – identification of UapA-TurboID interactome, post-ER interactions and Endocytic interactions

Volcano-plots were created using the VolcaNoseR2 online tool using the data processed in Perseus as previously described (Goedhart J. et al., 2020). Each plot summarizes only the significant differences [log2(fold-change)] in protein abundance in the x-axis and the – log10(p-value) of the Student‟s t-test performed from replicates in the y-axis. For UapA-TurboID and Δ3 PDB-LC-MS/MS experiments (**Figure 2A**), colored dots correspond to the upregulated proteins (blue) in UapA-TurboID strain in comparison with the negative control-Δ3 strain, and the gray to the unaffected proteins based on the cut-off of 0.5 for difference and p<0.05 for the significance of Student‟s t-test. For UapA-DYDY-TurboID and UapA-TurboID experiment (**Figure 3D**), blue colored dots indicate proteins upregulated in UapA-TurboID, red colored dots correspond to proteins upregulated in the UapA-DYDY-TurboID strain and gray colored unaffected proteins, respectively. For the particular figure, the cut-off of difference was set at 0.9 and the significance of Student‟s t-test at p<0.05. Lastly, for Δ*artA* UapA-TurboID and UapA-TurboID experiments (**Figure 4A**), the blue colored dots correspond to proteins enriched in wild type UapA-TurboID and green colored ones to those enriched in Δ*artA* UapA-TurboID strain, the cut-off of difference was set at 0.2 and the significance of Student‟s t-test at p<0.05.

Using the aforementioned cut-offs for each experiment we extracted the following lists, provided in Supplementary material: **UapA-TurboID interactome** (based on upregulated proteins (blue) in UapA-TurboID strain in comparison with the negative control-Δ3 strain, **Figure 2A**), putative **Post-ER interactions** (based on protein hits enriched in wild type UapA-TurboID in comparison with UapA-DYDY-TurboID, **Figure 3D**), and putative **Endocytic interactions** (based on proteins enriched in wild type UapA-TurboID in comparison with Δ*artA* UapA-TurboID strain, **Figure 4A**). Subsequently, for each Uniprot accession, we searched the best matched gene accession number at the FungiDB (https://fungidb.org accessed on 3 May 2023) using the BLAST search tool of the above database (Basenko et al., 2018; Camacho et al., 2009). In the aforementioned lists, any existing homologous genes from yeast genome DB (https://www.yeastgenome.org) are also incorporated (Cherry et al., 2012).

### 2.7 Protein Class Annotation and Trafficking factors identification

Protein classes were assigned to the identified proteins using PantherDB, a comprehensive protein classification system. The annotation process involved mapping the identified proteins in **UapA-TurboID interactome**, to known protein classes based on functional annotations and domain information. In **Figure 2B** only the most abundant subclasses are displayed, excluding the unclassified genes (799 out of 1296 identified in UapA-TurboID interactome, were annotated). The data for the protein classes, including the number of genes and the percentage, were extracted from Panther DB and the protein class distribution was visualized using a bar plot. Each protein class was depicted as a horizontal bar, with the length of the bar corresponding to the respective percentage. The number of genes for each protein class was displayed on top of each bar. The bar plot was generated using the Python programming language and the matplotlib library (version 3.9) (Hunter et al., 2007). Membrane traffic proteins (PC00150) concern 7.3% of the classified genes in UapA-TurboID interactome. Manual annotation was performed to identify the unclassified/not annotated Trafficking factors in the obtained datasets. Testing the presence of the trafficking factors on each dataset (**UapA-TurboID interactome, Post-ER interactions, Endocytic interactions**), we constructed the **Supplementary Table S1, S2, S3, S4**, where the -log(p-value) and the log2(fold-change) for each protein hit is displayed. Note that Supplementary Table S2 shows the same proteins as Supplementary S1 but with -log(p-value) and the log2(fold-change) from UapA-DYDY-TurboID and UapA-TurboID PDB-LC-MS/MS results.

### 2.8 Gene Enrichment Analysis

Gene enrichment analysis was performed to gain insights into the functional implications of the identified proteins. ShinyGo (version 0.77), a gene enrichment analysis tool, was employed with the experimental background for each pull-down experiment (the background of each experiment is presented in the **Supplementary: Background** list) (Xijin et al., 2019). The analysis involved identifying overrepresented gene ontology terms and pathways, using GO: Biological process and GO: Cellular component databases for *A. nidulans*, among the Supplementary material: **UapA-TurboID interactome, Post-ER interactions, Endocytic interactions**. FDR is calculated based on nominal P-value from the hypergeometric test. Fold Enrichment is defined as the percentage of genes in a given list belonging to a pathway, divided by the corresponding percentage in the background. The FDR cutoff was set at 0.05 for UapA-TurboID interactome and Post-ER interactions, and 0.2 for Endocytic interactions. The enriched pathways were selected by FDR, sorted by Fold Enrichment. The results of the gene enrichment analysis for Gene Ontology (GO) Biological Process and (GO) Cellular Component terms retrieved from ShinyGO, were visualized using Python-matplotlib to create the dot plot representing the enriched pathways in **Figure 2C**-**D**, **Figure 3F**, **Figure 4B**.

### 2.9 Heatmaps

Heatmaps for each PDB-LC-MS/MS experiment were generated using Perseus software. The data was preprocessed and imputed for missing values. Euclidean clustering was performed on the columns using Z-score transformation. The resulting heat maps provided visual representation of protein abundance across samples.

### 2.10 Transport Assays

Kinetic analysis of UapA in the TNO2A7 and mutant strains (*glo3*Δ, *gcs1*Δ) was measured by estimating uptake rates of [^3^H]-xanthine uptake (40 Ci. mmol^-1^, Moravek Biochemicals, CA, USA), as previously described in Krypotou and Diallinas (2014). In brief, [3 H]-xanthine uptake was assayed in *A. nidulans* conidiospores germinating for 4 h at 37° C at 140 rpm, in liquid MM, pH 6.8. Initial velocities were measured on 10^7^ conidiospores/100 μL by incubation with concentration of 0.3 μM of [^3^H]-xanthine at 37° C. All transport assays were carried out at least in 2 independent experiments and the measurements in triplicate. Standard deviation was <20%. Results were analyzed in Microsoft Excel. The raw data extracted were visualized in a bar plot representing the % [^3^H]-xanthine uptake rate for each strain as percentages of initial uptake rates compared to the wt UapA rate. UapA initial uptake rate in the isogenic TNO2A7 is arbitrarily taken as 100%. The plot was created in Python, using the Matplotlib library.

### 2.11 Fluorescence microscopy and statistical analysis

Samples were prepared as previously described (Martzoukou et al., 2017). Unless otherwise stated, conidiospores were incubated overnight in glass bottom 35-mm l-dishes (Ibidi, Lab Supplies Scientific SA, Hellas) in liquid minimal media, for 16–22 h at 25°C, under conditions of transcriptional repression of the studied membrane cargoes, UapA and SynA (Dimou et al., 2020). For following the subcellular trafficking of neosynthesized UapA-GFP, we used either the *uapA* native promoter or the regulatable *alcA* promoter. The *uapA* promoter can be tightly repressed in the presence of 10 mM ammonium tartrate supplied as a nitrogen source in the growth medium, while the *alcA* promoter was repressed in the presence of 1% (w/v) glucose as a carbon source. The *alcA* promoter was also used for following the trafficking of the apical membrane cargo SynA, tagged with GFP or mCherry. Genes involved in trafficking, expressed under the *thiA_p_* promoter, i.e. *thiA_p_-copA* and *thiA_p_-arfA*, were transcriptionally repressed in the presence of 10 μM thiamine in the growth media. As addition of thiamine *ab initio* in the culture led to total arrest of spore germination and germling formation, an 8 h time window without thiamine at the start of spore incubation was introduced, in order to let conidiospores germinate until the stage of young germlings. Then thiamine was added in the media (16-18 h) under conditions where cargo (UapA, SynA) expression was repressed. Transcriptional repression of the studied membrane cargoes (UapA, SynA) was followed by a derepression period, through a shift in media containing either NaNO3 as a nitrogen source for the native *uapA* promoter, or 0.1% (w/v) fructose as a carbon source for the *alcA* promoter.

Images were obtained using an inverted Zeiss Axio Observer Z1 equipped with an AxioCam HR R3 camera. Contrast adjustment, area selection and color combining were made using the Zen lite 2012 software. Scale bars were added using the FigureJ plugin of the ImageJ software. Images were further processed and annotated in Adobe Photoshop CS4 Extended version 11.0.2. Technical replicates correspond to different hyphal cells observed within each sample, while biological replicates correspond to different samples (Martzoukou et al., 2017). For quantifying co-localization, Pearson‟s correlation coefficient (PCC) above thresholds, for a selected region of interest (ROI), was calculated using the ICY co-localization studio plugin (pixel-based method) (http://icy.bioimageanalysis.org/). The fluorescence intensity of UapA-GFP in wt and Δ*glo3* was quantified using the ICY software as described in Dimou et al., 2020. In brief, two ROIs in the same region were drawn manually, using the Area Selection tool, one including both the PM and the cytoplasm and another identical one, excluding the PM. The PM/cytoplasmic mean fluorescence intensity ratios for each strain are presented in box scatter plots in **Figure7C,** using the GraphPad Prism software. To test the significance of differences in PM/cytoplasmic fluorescence of measurements between the two strains, an unpaired (two-sample) t-test was performed using the GraphPad Prism software. Confidence interval was set to 95%.

### 2.12 Protein extraction and western blotting

Total protein extraction was performed as previously described by Dimou et al., 2020. For strains shown in **Figure 1D** and **Figure 3D**, dry mycelia from cultures grown in minimal media supplemented with 10 mM NH_4_^+^, 10 mM NaNO_3_ or 0.5 mM uric acid at 30°C for 16 h were used. Biotin was added to a final concentration of 50 μM. Total proteins (50 μg, estimated by Bradford assays) were separated in a 10 % (w/v) polyacrylamide gel and then transferred on PVDF membranes (GE Healthcare Life Sciences, Amersham). Immunodetection was performed with an anti-HA monoclonal antibody [HA Epitope Tag (YPYDVPDYA) Mouse Monoclonal Antibody–AM20783PU-N, OriGene], an HRP-linked antibody (7076, Cell Signaling Technology Inc.) and Streptavidin-HRP conjugate (Cytiva). Colloidal Coomassie G-250 Staining for Proteins (CBB) was employed as described in Dyballa and Metzger, 2009. Ponceau S (2%) stain was employed as a loading control in **Figure3.C** (Sander et al., 2019).

**Figure 1.**
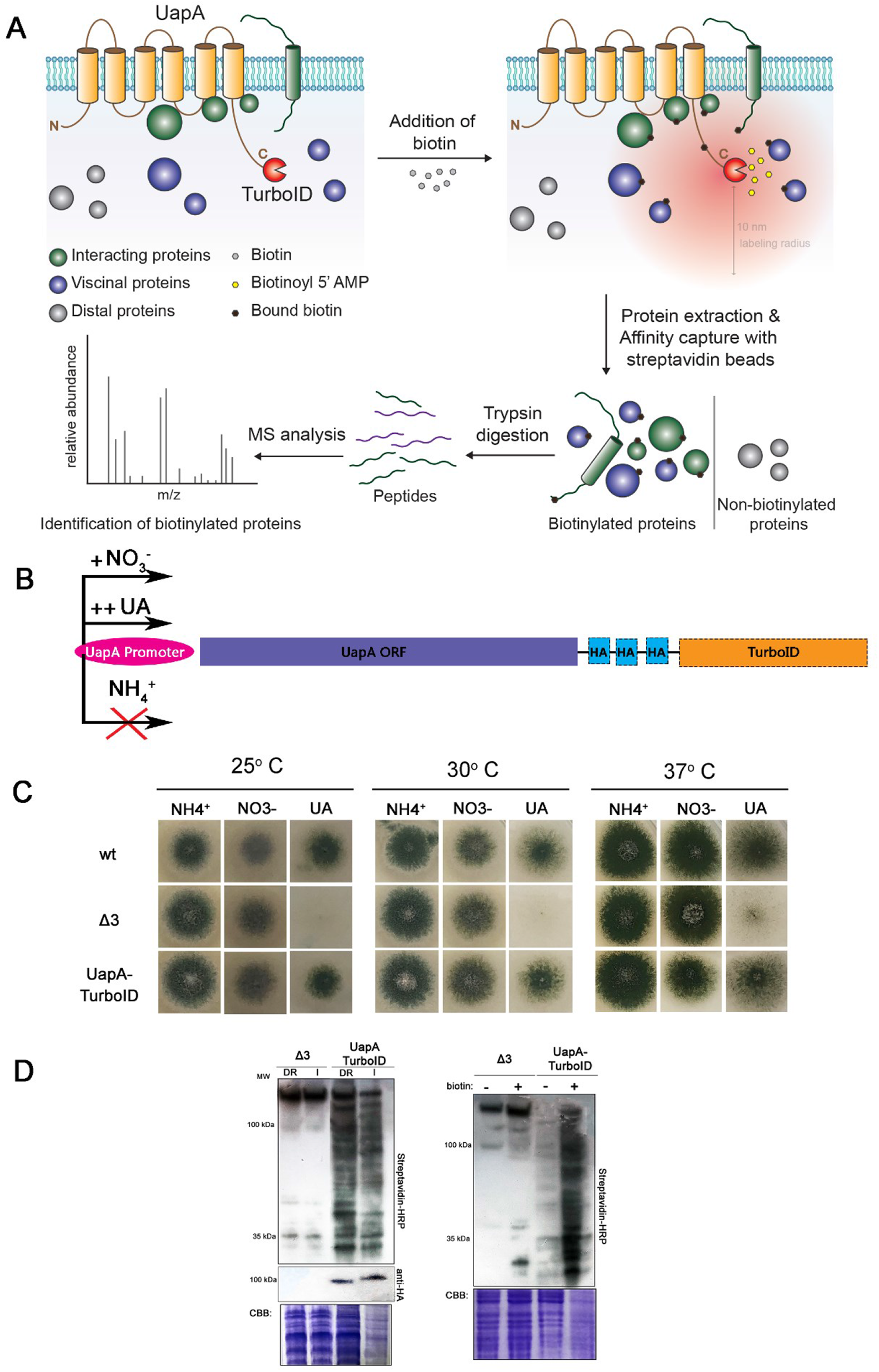
Development of a system to identify UapA interactors via PDB. **A.** Cartoon illustrating the proximity-dependent biotinylation (PDB) of UapA-TurboID and the experimental workflow of PDB-LC-MS/MS analysis. **B.** UapA-TurboID chimeric construct used in this study. The expression of UapA-TurboID is tightly regulated by the *uapA* native promoter. Namely, its expression is repressed in the presence of ammonium (NH_4_^+^), derepressed in nitrate (NO_3_^-^) and induced in uric acid (UA). Thus, the chimeric UapA-TurboID expression is dependent on the nitrogen source of the culture media. **C.** Growth test of the strain expressing UapA-TurboID on selected N sources at 25, 30 and 37°C, compared to a wild-type strain (wt) and a Δ*uapA*, Δ*uapC* and Δ*azgA* negative control strain (Δ3). N sources shown are: 10 mM ammonium tartrate (NH_4_^+^), 10 mM sodium nitrate (NO_3_^-^) and 0.5 mM uric acid (UA). Notice that the chimeric UapA-TurboID is functional, as the relative strain can grow on minimal media containing UA as the sole N source. **D.** Left panel: Western blot analysis with streptavidin-HRP showing UapA-TurboID potential to label protein interactors under conditions where the *uapA* promoter is derepressed (DR) or induced (I). Strains were grown under derepressing (NO_3_^-^) or inducing (UA) conditions upon biotin treatment. The Δ3 strain was used as a control to visualize the presence of endogenously biotinylated proteins. The Coomassie Brilliant Blue-stained membrane (CBB) is shown as loading control. Right panel: Similar western blot analysis showing the dependency of UapA-TurboID activity on biotin addition under derepressing conditions. Addition of external biotin increases dramatically the activity of UapA-TurboID.

**Figure 2.**
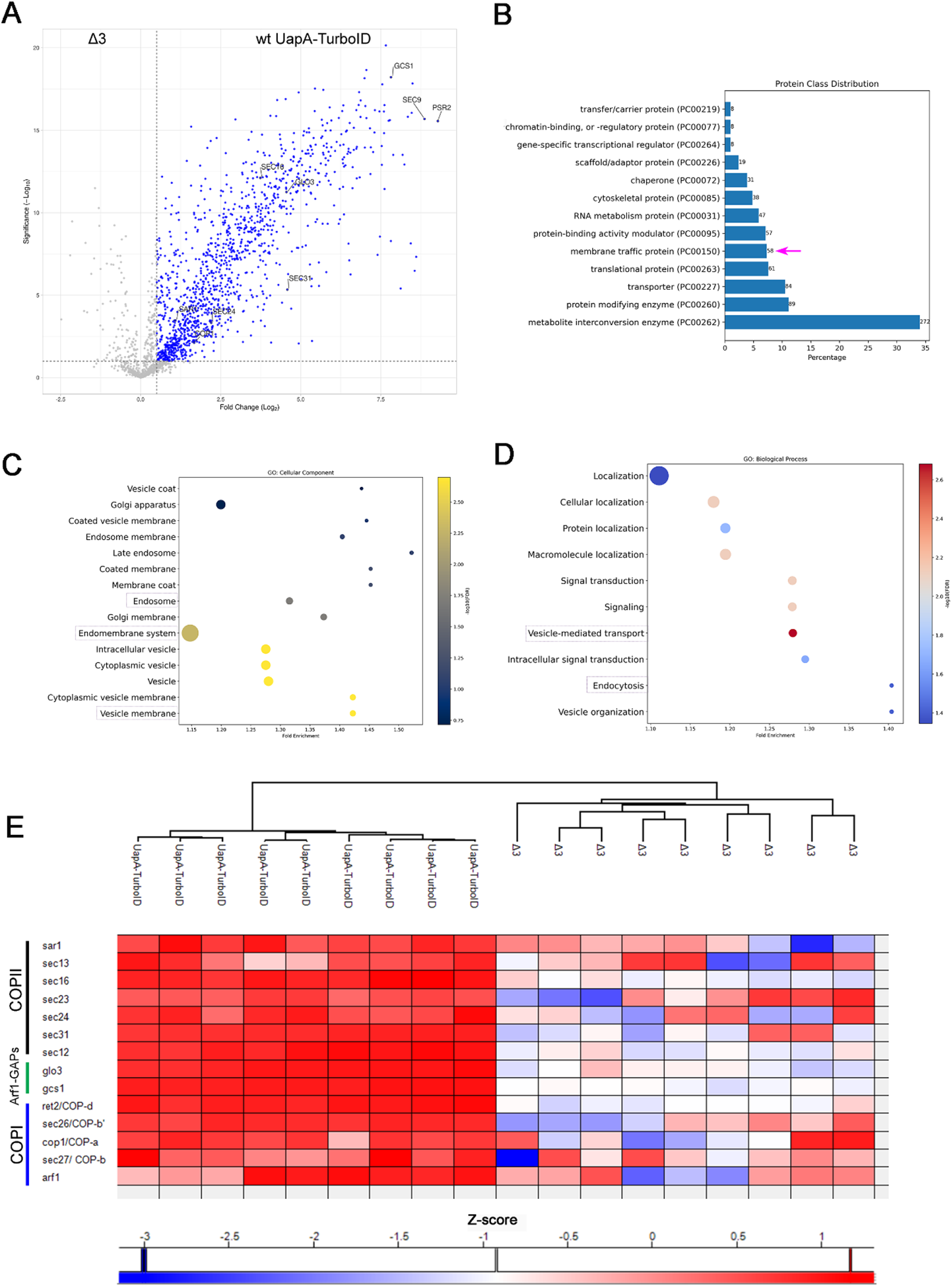
The interactome of wt UapA. **A.** Volcano plot of PDB LC-MS/MS comparing UapA-TurboID with the isogenic negative control Δ3 strain. Blue colored dots correspond to UapA-TurboID interactome, according to 0.5 cut-off for log_2_ (Fold Change) and p<0.05 for the significance of Student‟s t-test. **B.** Protein class annotation of the identified UapA-TurboID interactome in PantherDB. Only the most abundant subclasses are displayed, excluding the unclassified genes. Concerning the genes of interest, 7.3% (58 genes) of the UapA-TurboID interactome includes proteins related to membrane cargo traffic (PC00150). **C.** Bar plot showing GO-TERM enriched cellular components in UapA-TurboID interactome. Dots size is indicative of the number of genes assigned to a given GO-term, color bar depicts the log_10_FDR and fold enrichment is shown. GO-terms concerning cellular components of interest are noted. **D.** Bar plot presenting GO-TERM enriched biological processes in UapA-TurboID interactome. Dots size is indicative of the number of genes assigned to a given GO-term, color bar depicts the log_10_FDR and fold enrichment is shown. GO-terms concerning biological processes relevant to membrane trafficking are noted. **E.** Heat map highlighting the significant enrichment of COPII and COPI components, and two selected Arf1 GAPs, Glo3 and Gcs1, in the UapA-TurboID interactome. The color bar indicates the z-score for the protein hits presented on the heat map.

**Figure 3.**
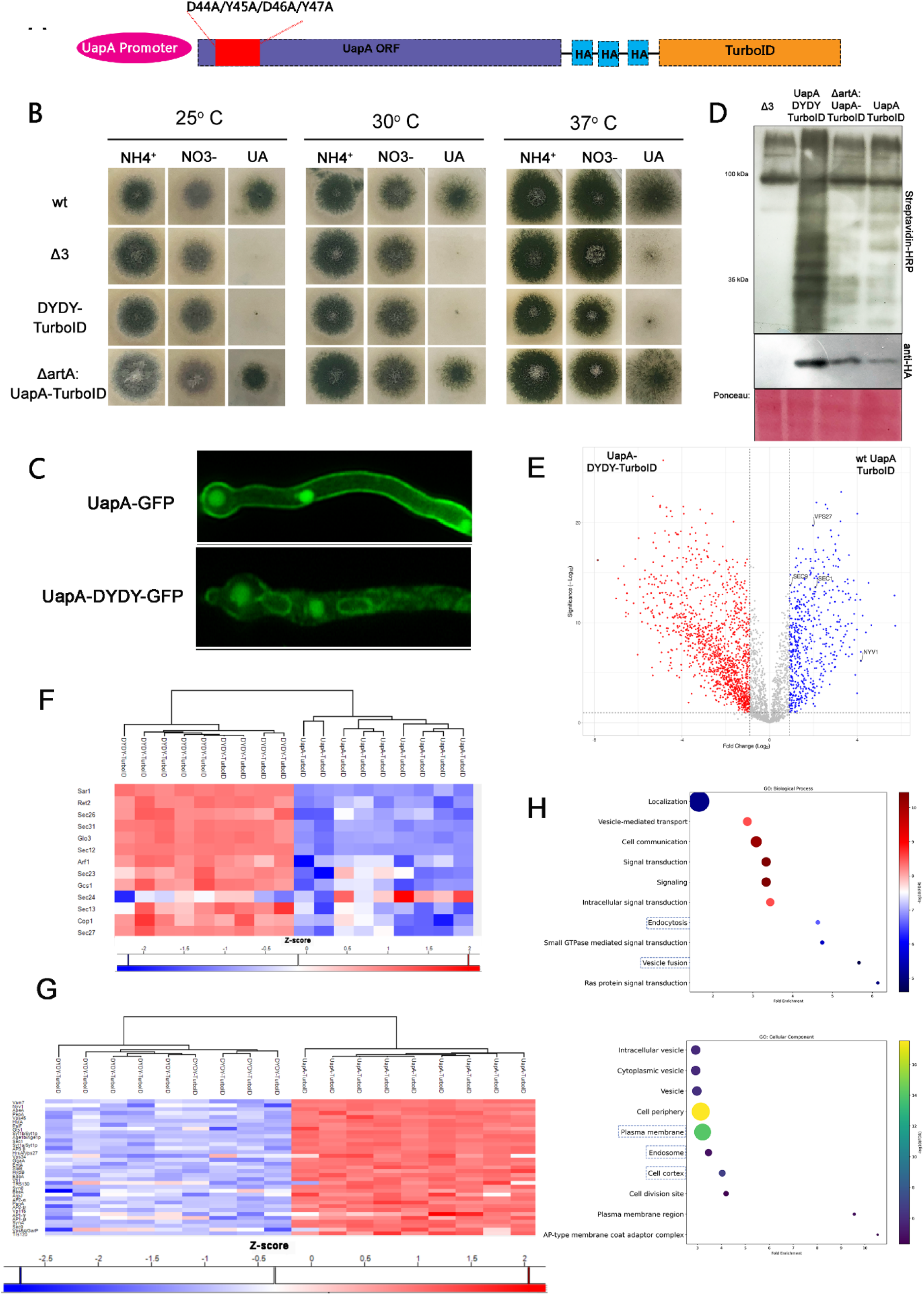
PDB assays in the UapA-DYDY mutant impaired in ER-exit. **A.** UapA-DYDY-TurboID chimeric construct. In this mutant version of UapA, a quadruple alanine substitution was introduced to the conserved motif Asp-Tyr-Asp-Tyr (DYDY) located in the cytosolic N-terminal region of the transporter. **B.** Growth tests of UapA-DYDY-TurboID on selected N sources at 25, 30 and 37°C, compared to a wt strain and a Δ*uapA*, Δ*uapC* and Δ*azgA* strain (Δ3). A strain expressing UapA-TurboID in an endocytosis defective background (Δ*artA* UapA-TurboID) is also shown. N sources tested are: 10 mM ammonium tartrate (NH_4_^+^), 10 mM sodium nitrate (NO_3_^-^) and 0.5 mM uric acid (UA). UapA-DYDY-TurboID lacks transport activity and thus cannot grow on uric acid. Δ*artA* UapA-TurboID can grow normally on minimal media containing uric acid as the sole N source. **C.** Epifluorescence microscopy showing the subcellular localization of UapA-DYDY-GFP mutant and wt UapA-GFP in the presence of NO_3_^-^ as nitrogen source. Notice the perinuclear localization (rings) of the mutant indicative of ER retention. **D.** Western blot analysis using streptavidin-HRP and anti-HA of protein extracts from UapA-TurboID, UapA-DYDY-TurboID and Δ*artA* UapA-TurboID strains, grown in derepressing (NO_3_^-^) conditions along with biotin addition. The Ponceau-stained membrane is shown as loading control. **E.** Volcano plot of PDB LC-MS/MS comparing UapA-TurboID (right) with UapA-DYDY-TurboID strain (left). Blue colored dots indicate enriched proteins in wild type UapA-TurboID and red colored dots correspond to proteins enriched in the UapA-DYDY-TurboID according to 0.9 cut-off for log_2_ (Fold Change) and p<0.05 for the significance of Student‟s t-test. From this experiment, protein hits enriched in wt UapA-TurboID (blue colored dots) indicate post-ER interactions of UapA. **F.** Heat map depicting the differential enrichment of COPII, COPI components and two selected Arf1 GAPs, Glo3 and Gcs1, in UapA-DYDY-TurboID and in wild type UapA-TurboID. Notably, all protein hits presented in the heat map are enriched in UapA-DYDY, with the exception of Sec24. Thus, Sec24 is the only COPII component mainly enriched in the wt rather than in the ER-retained UapA. **G.** Heat map presenting trafficking factors found enriched in wild type UapA-TurboID, thus consisting of putative post-ER exit interactions. **H.** Bar plots displaying GO-TERM enriched biological processes and cellular components in protein hits enriched in UapA-TurboID (blue dots Figure 3E, Supplementary Table S2). Dot size is indicative of the number of genes assigned to a given GO-term, color bar depicts the log_10_FDR and fold enrichment is shown. GO-terms concerning cellular components of interest are noted. As can be seen, cellular components such as plasma membrane, cell cortex, cell periphery and endosome are enriched in the post-ER dataset, indicating that comparison between the wild type and the ER-retained mutant can possible reveal interactions occurring in later steps of vesicular secretion.

To test the efficacy of thiamine repression in *copA* and *arfA* (**Figures 5B** and **6B**), cultures were grown on minimal media (10 mM NH_4_^+^) without thiamine for 8 h, following by supplementation of 10 μΜ thiamine for additional 16 h incubation at 25°C. Mycelia were then collected. Protein extraction, separation and electroblotting were carried out as mentioned previously. Proteins were detected by a standard Western blot analysis using either an anti-FLAG antibody (Agrisera, AAS15 3037) for *thiA_p_*-^FLAG^*copA* or an anti-GFP antibody (11814460001, Roche Diagnostics) for *thiA_p_-arfA*-GFP and an anti-actin monoclonal (C4) antibody (SKU0869100-CF, MP Biomedicals, Europe). Blots were developed using the Lumi Sensor Chemiluminescent HRP Substrate kit (Genscript, United States) and SuperRX Fuji medical X-Ray films (Fuji FILM, Europe).

**Figure 4.**
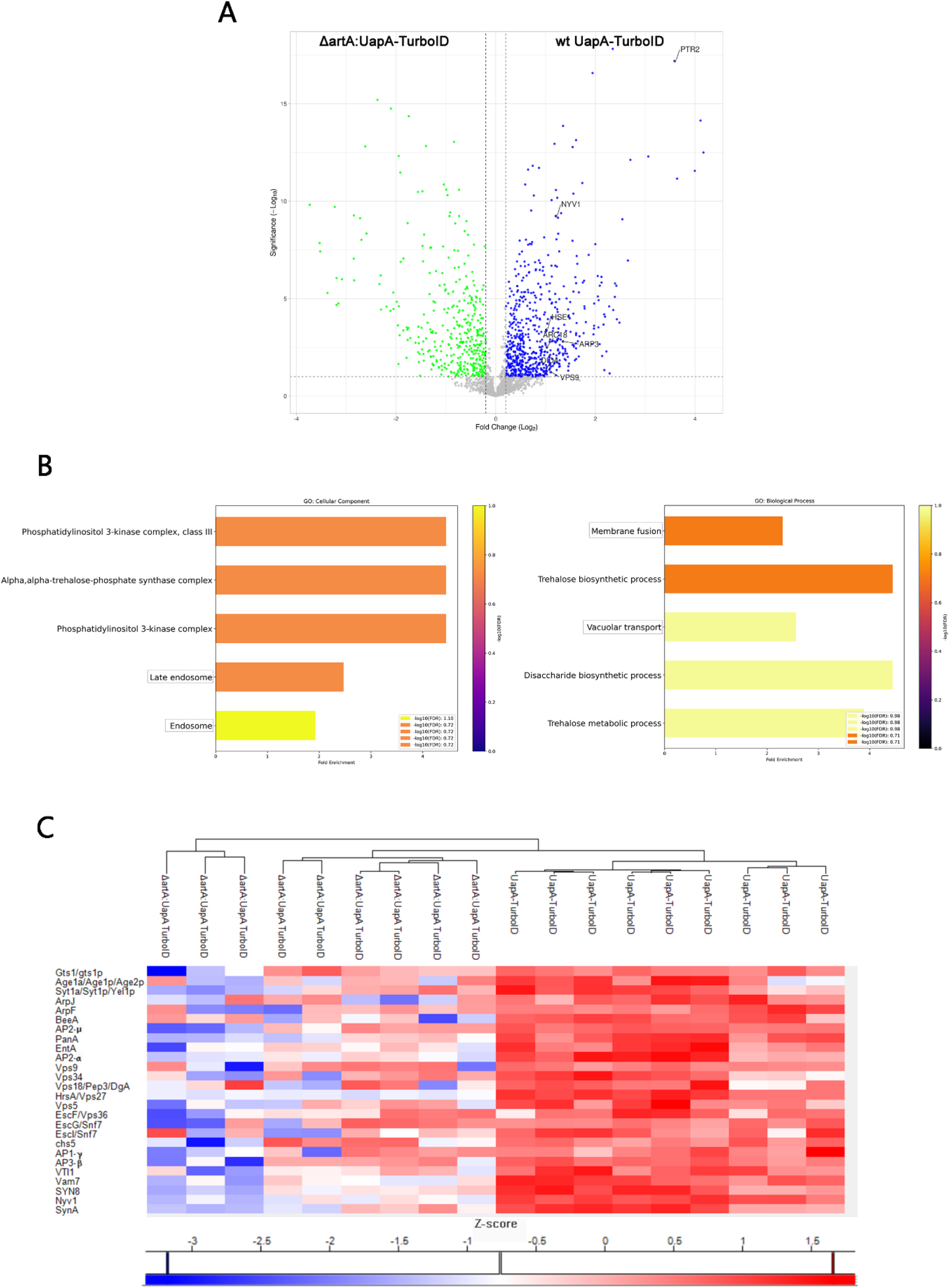
The interactome of Δ*artA* UapA-TurboID. **A.** Volcano plot of PDB LC-MS/MS comparing UapA-TurboID with Δ*artA* UapA-TurboID strain. Blue colored dots correspond to proteins enriched in wild type UapA-TurboID and green colored ones to those enriched in Δ*artA* UapA-TurboID strain, according to 0.2 cut-off for log_2_ (Fold Change) and p<0.05 for the significance of Student‟s t-test. From the particular experiment, proteins enriched in the wild type UapA-TurboID strain suggest putative endocytic interactions of UapA. **B.** Bar plots displaying GO-TERM enriched biological processes and cellular components in protein hits enriched in wild type UapA-TurboID (blue dots Figure 4A, Supplementary Table S6, S7). Dot size is indicative of the number of genes assigned to a given GO-term, color bar depicts the log_10_FDR and fold enrichment is shown. In the bar plot, biological processes concerning vacuolar transport, membrane fission and cellular components, such as endosomes or late endosomes, are pointed out. The gene enrichment analysis verified that the comparison between the wild type and Δ*artA* UapA-TurboID can reveal putative endocytic interactions, as postulated. **C.** Heat map presenting trafficking factors found enriched in wild type UapA-TurboID relative to Δ*artA* UapA-TurboID (blue dots Figure 4A), thus consisting of putative endocytic interactions.

**Figure 5.**
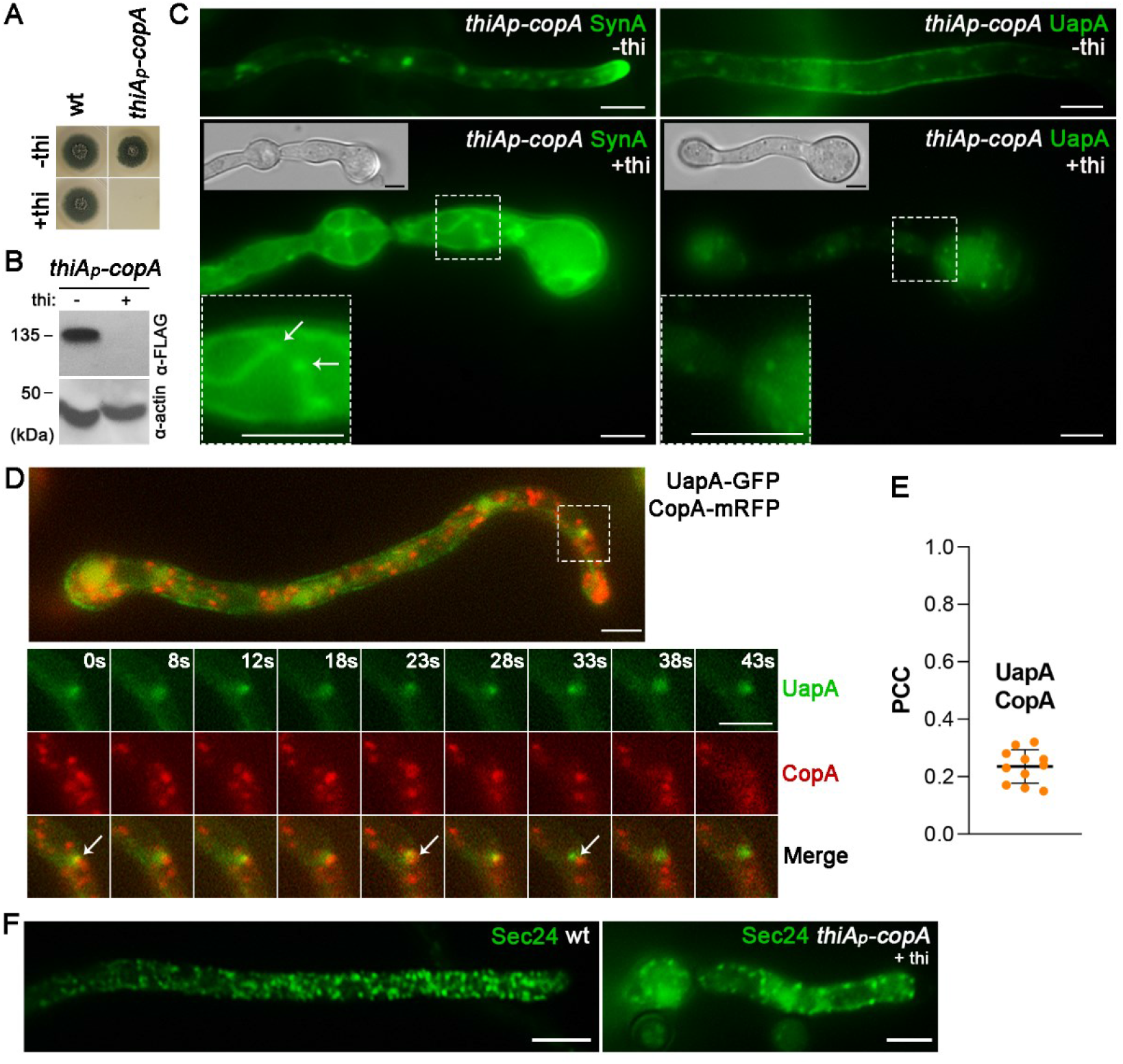
CopA is essential for anterograde cargo trafficking, germination and growth. **A.** Growth test analysis of the *thiA_p_-copA* strain compared to a wild-type control strain in the presence (+thi) or absence (-thi) of thiamine from the media, showing that CopA repression impedes colony formation. **B.** In the absence of thiamine (-thi) from the growth medium, CopA is expressed, while upon addition of thiamine (+thi), the expression of CopA is tightly repressed. Proteins are detected by a standard Western blot analysis using the anti-FLAG antibody. Equal loading and protein steady-state levels are normalized against the amount of actin, detected with an anti-actin antibody. **C.** Epifluorescence microscopy analysis of the subcellular localization of GFP-SynA and UapA-GFP under conditions where *copA* transcription is derepressed (-thi) or repressed by addition of thiamine (+thi). *De novo* synthesis of cargoes takes place after full repression of CopA is achieved (>16 h). Repression of *copA* abolishes the proper localization of SynA to the PM of the hyphal apex and leads to ER-retention (white arrows). Under the same repressing conditions, UapA PM localization is lost. Scale bars: 5 μM. **D.** Co-localization analysis and relevant quantification of a strain co-expressing *de novo* made UapA-GFP with CopA-mRFP. Quantification by calculating Pearson‟s correlation coefficient (PCC) shows no co-localization of UapA with CopA (PCC = 0.24 ± 0.06). Biological/technical replicates: 2/11. Time-series experiment showing a transient colocalization of UapA with CopA for less than 1 min (0-28s). Scale bar: 5 μM **E.** Epifluorescence microscopy analysis of the subcellular localization of Sec24-GFP in a *thiAp-copA* genetic background. Notice that repressing CopA expression (+thi) did not affect the formation of Sec24-labeled fluorescent foci. Scale bar: 5 μm.

**Figure 6.**
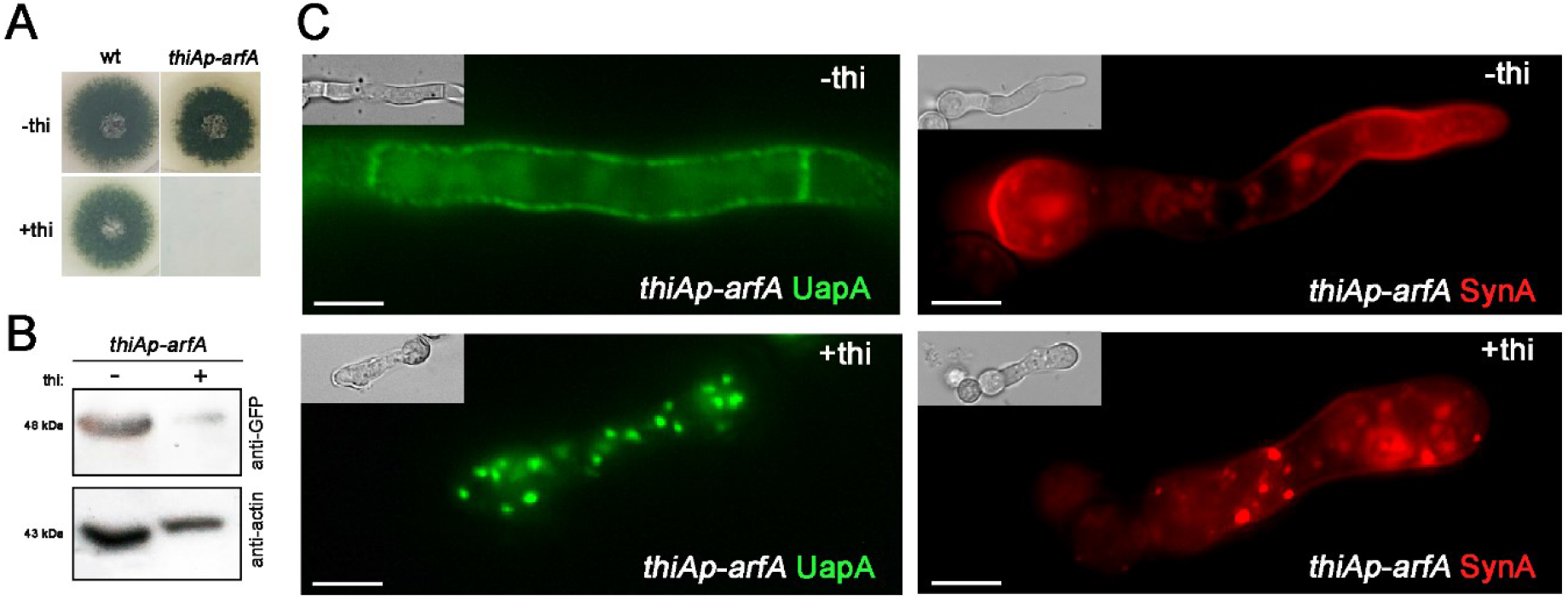
ArfA is essential for anterograde cargo trafficking, germination and growth. **A.** Growth test showing that ArfA expression is essential for growth as its transcriptional repression by thiamine (+thi) leads to the absence of colony formation. **B.** Western blot analysis using an anti-GFP antibody to detect ArfA-GFP in a strain where the endogenous *arfA* promoter has been replaced by the *thiA* promoter. Equal loading and protein steady-state levels are normalized against the amount of actin, detected with an anti-actin antibody. In the presence of thiamine (+thi) in the culture media, ArfA-GFP (∼48 kDa) is almost undetectable. **C.** Epifluorescence microscopy analysis of the subcellular localization of UapA and SynA under conditions where *arfA* transcription is derepressed (-thi) or repressed by addition of thiamine (+thi). Repression of *arfA* abolishes the proper localization of both cargoes to the PM, which appear in prominent cytosolic aggregates. Scale bars: 5 μm.

## 3. Results and Discussion

### 3.1 Development of a system to identify UapA interactors via PDB

Our goal was to use proximity dependent biotinylation (PDB) assays coupled with LC-MS/MS to identify transient interactors of the UapA xanthine-uric acid transporter (Diallinas 2016) during its biogenesis, that is, from the moment of its *de novo* synthesis in the ER to its steady state localization to the PM (Diallinas and Martzoukou, 2019). In contrast to many other affinity capture approaches for studying protein-protein interactions, PDB does not rely on physical protein-protein binding, allowing the identification of protein proximities of weak or transient and dynamic nature. For the PDB assays we used TurboID, an engineered biotin ligase that uses ATP to convert biotin into biotin-AMP, a reactive intermediate that covalently labels proximal proteins. TurboID has been reported to have higher activity than other biotin ligase–related proximity labeling methods, such as BioID, enabling higher temporal resolution and broader application *in vivo* (Cho et al., 2020). The rationale of our approach is depicted in **Figure 1A**.

For identifying proteins interacting with the UapA transporter, we constructed a *uapA-TurboID* fusion DNA cassette (**Figure 1B**). The *uapA-TurboID* fusion cassette was used to transform a recipient strain that genetically lacks the endogenous *uapA* gene, as well as, the other two genes encoding major purine transporters, namely AzgA and UapC (Δ3 strain; Pantazopoulou et al., 2007). This genetic background permits the direct functional validation of UapA-TurboID chimeras by simple growth tests on uric acid as a N source. The UapA-TurboID fusion was designed to be expressed by the native promoter of *uapA*, which allows tight transcriptional control of expression in response to the N source supplied in the growth medium (see also later).

**Figure 1C** shows that a strain expressing UapA-TurboID could grow on uric acid as a sole N source, confirming that UapA-TurboID was functional in respect to transport activity. To test whether the selected strain expressing UapA-TurboID possess TurboID-dependent biotin ligase activity, we performed western blot analysis using Streptavidin-HRP antibody against total protein extracts obtained under different conditions of UapA-TurboID expression (**Figure 1D**). Transcription from the *uapA* native promoter is repressed by ammonium, derepressed by other nitrogen sources (e.g. nitrate), or induced in the presence of substrates of UapA (e.g., uric acid). Therefore, cultures were grown in derepressed (nitrate) or induced (uric acid) conditions for 16h, with or without external biotin supplementation prior to protein isolation. As a negative control, we also included the isogenic Δ3 strain, lacking the UapA-TurboID fusion. The control strain was expected to reveal the profile of endogenously biotinylated proteins, as fungi are known to possess endogenous biotin and biotin ligases (Fenech et al., 2023). **Figure 1D** shows that UapA-TurboID mediates biotinylation *in vivo* when its expression is transcriptionally derepressed or induced. In addition, the presence of external biotin led to an increase in biotinylated proteins, as assessed under derepressed conditions. The negative control showed a minimal level of endogenously biotinylated proteins that depended little on the conditions affecting UapA-TurboID expression or addition of biotin. Based on these results, we chose to use growth conditions that allow derepressed levels of UapA-TurboID transcription, in the presence of externally supplied biotin, for downstream PDB assays.

### 3.2 The interactome of wt UapA

We chose to use conditions that lead to derepression (nitrate as N source) rather than induction (uric acid as N source) of UapA-TurboID transcription, despite the fact that the latter lead to higher levels of biotinylated proteins, because the presence of uric acid also leads to very high rates of UapA endocytic degradation (Gournas et al., 2010; Karachaliou et al., 2013). This would lead to a more complicated interactome of UapA, including proteins involved in ubiquitination, internalization and endosomal trafficking to vacuoles, which is beyond our principal scope to identify interactors of UapA involved in anterograde translocation to the PM. Furthermore, we chose to reduce the time frame of derepression of UapA-TurboID expression to 4h in order to avoid over-accumulation of UapA molecules in the PM, which might „mask‟ interactions occurring at earlier steps of UapA biogenesis. Furthermore, reducing the time of derepression also leads to reduction of the low level turnover occurring after over-accumulation of UapA in the PM. In brief, proteins were extracted from cultures grown overnight (16h) in the presence of ammonium (repressed conditions) and shifted for 4h in nitrate media (derepressed conditions) supplemented with 50 μΜ biotin. For isolating biotinylated proteins, we performed streptavidin pull-downs coupled with liquid chromatography and tandem mass spectrometry (LC-MS/MS) analysis.

To distinguish between physiological/functional relevant interactions from those identified due to stochastic close proximity, prior biotinylation, or nonspecific binding, we included the negative control strain Δ3 in our experimental setup. To that end, the biotinylated proteome of UapA-TurboID was compared to the equivalent proteome of the negative control strain, focusing on profound changes in protein abundance. The analysis revealed 1296 proteins as potential UapA-interacting partners based on differences in abundance [log_2_(fold change)]. More specifically, value differences above 0.5 were examined and listed in **Supplementary Table S1. Figure 2A** shows the volcano plot depicting proteins enriched in UapA-TurboID relative to the control stain Δ3. The acquired UapA-TurboID interactome was then classified in terms of protein classes using PANTHERdb (**Figure 2B**) (Thomas, P. D. et al., 2021). The result validates that the experimental workflow adapted herein was able to identify annotated membrane traffic proteins (PC00150). Subsequently, gene enrichment analysis was performed to gain insights into the functional implications of the identified interactome. **Figures 2C** and **2D** depict the results of the GO-term enrichment analysis in respect to subcellular compartment and biological process respectively. As can be seen, GOs concerning vesicle-mediated transport, endocytosis, and sorting to vacuoles are all enriched, as anticipated for a plasma membrane transporter. Selected pathways relative to membrane trafficking are highlighted (**Figure 2C** and **2D**). Notice that in the present work, we use the protein nomenclature of *S. cerevisiae*, unless a protein has been given a name in *A. nidulans*.

For an in depth characterization of the proteins mediating vesicular secretion, we manually annotated trafficking factors present in the acquired interactome and compiled them in **Supplementary Table S2.** 70 proteins known to be involved in membrane cargo trafficking were identified in the UapA-TurboID interactome. Among those, all COPII coat proteins (Sar1, Sec24, Sec23, Sec31, Sec13, Sec16 and Sec12) and several COPI components (CopA^Cop1,^ Sec26, Sec27 and Ret22) were included. Prominent scores were obtained with proteins regulating ArfA^Arf1^, the major small GTPase regulating intra-Golgi transport (Lee and Shaw, 2008). Specifically, guanine nucleotide exchange factors (GEFs) Syt1a/b and HypB^Sec7^ and guanine activating proteins (GAPs) Glo3, Gcs1 and Age1b (Poon et al., 1999; Zhang et al., 2003, Hernandez-Gonzalez, 2018) were highly enriched. A heat map highlighting the significant increase of COPII, COPI and other trafficking proteins on UapA- urboID interactome is presented in **Figure 2E**. The importance of the interactions with COPI becomes apparent later.

In general, proteins involved in different steps of vesicular secretion were isolated using the current methodology. Specifically, SNARE proteins involved in membrane fusion, such as Sec22, BetA, SedV, Vam7, SynA, Syn8 and Sec9, were significantly enriched. Among Rab GTPases, RabE and RabS, were included in the obtained interactome. Several factors involved in post-Golgi trafficking (e.g., AP-1 and AP-3 subunits, GgaA, ClaL; ref), proteins associated with ER-PM contacts (e.g., Tcb1-Tcb3; ref), ubiquitination (e.g., HulA^Rsp5^ ref), endocytosis (e.g., AP-2 subunits, SlaB; ref), vacuolar sorting (e.g., ESCRTs; ref) and exocyst core components (e.g., Sec5, Sec10, Sec15) or effectors (e.g., Sec1), were also highly enriched.

In summary, we obtained putative UapA interactors localized all along the conventional trafficking pathway (i.e., ERes/COPII, COPI, early and late Golgi, post-Golgi, exocyst, endocytosis, vacuolar sorting). However, based on our previous systematic genetic approaches (Dimou et al., 2020), we concluded that several of these interactions should not be physiologically relevant, and are probably the result of very close proximity of subcellular compartments involved in cargo trafficking, such as the ER and the Golgi, but also the cortical ER and the PM. This conclusion is based on strong direct genetic evidence showing that UapA traffics to the PM via Golgi bypass. In particular, it has been shown that genetic knockout or knockdown of several Golgi-related proteins, which appear here as UapA interactors (e.g., SedV, HypB, RabE, AP-1, AP-3), are redundant for UapA localization to the PM (Martzoukou et al., 2018, Dimou et al., 2020). Thus, PDB assays led to an apparent paradoxical result, which in turn strongly suggests that our system inevitably lacks specificity. This however does not exclude that several of the interactions identified might well be physiologically relevant for UapA localization to the PM. Thus, the challenge was to reduce the background „noise‟ of our system, before prioritizing and validating a number of putative UapA interactions. To this direction, we proceeded in obtaining the interactome of a UapA mutant version trapped in the ER (UapA-DYDY-TurboID) or the interactome of wt UapA expressed in a genetic background where endocytosis is compromised (Δ*artA* UapA-TurboID).

### 3.3 The UapA interactome of mutants impaired in ER-exit or endocytosis

Mutations in the conserved motif Asp-Tyr-Asp-Tyr (DYDY) located in the cytosolic N-terminal region of UapA have been shown to lead to ER-retention and lack of transport activity, despite evidence that the transporter can still bind its substrates with normal affinity (Martzoukou et al., 2015). This is best exemplified by a strain expressing a version of UapA where DYDY is replaced by alanine. This mutant lacks transport activity and thus cannot grow on uric acid because UapA is retained in the ER. Given this mutant UapA version fused with GFP marks both the nuclear and cortical ER suggests that UapA is not localized at ER-exit sites (ERes), but partitions and remains rather stable in the entire ER network. In fact, the DYDY motif is thought to be an element necessary for recognition by Sec24/Sec23 at ERes and thus necessary for UapA packing into COPII vesicles and ER-exit. Thus, this mutant version of UapA is expected to lose all trafficking-related protein interactions, which in the wild type UapA take place at ERes and during COPII vesicle formation. For that reason, we thought of using the UapA-DYDY/AAAA mutant in PDB assays and comparing its interactome with that of the wild type UapA. This would in principle allow us to identify true interactions of UapA taking place after ER-exit, keeping in mind possible exceptions where the ER is in very close contact with other compartments, as for example the early-Golgi or contacts of cortical ER with PM.

To that end, we constructed and functionally analyzed a strain expressing UapA-DYDY-TurboID (simplified naming of UapA-DYDY/AAAA-TurboID), as described for wild type UapA-TurboID (**Figure 3A**). As expected, this strain could not grow on uric acid (i.e., lacked UapA transport activity) and showed retention of UapA in the ER (**Figure 3B** and **3C**). Subsequently, we confirmed by western blot analysis using streptavidin-HRP antibody that UapA-DYDY-TurboID can be used for *in vivo* proximity labeling (**Figure 3D**), and then employed PDB assays as described for wild type UapA.

The comparison of the interactome of UapA-DYDY-TurboID with the wild type UapA-TurboID revealed 245 protein hits enriched in the wild type UapA relative to UapA-DYDY, with value difference above 0.9 [log_2_(fold change)]. These protein interactions lost in UapA-DYDY were considered to occur specifically at ERes or after ER-exit. These „post-ER‟ interactions are displayed on the volcano plot of **Figure 3E** (blue colored dots) and listed in **Supplementary Table S3**. To further characterize the aforementioned dataset, GO-term enrichment analysis was employed in respect to cellular components and biological processes. The result shows that cellular components such as PM, cell cortex, cell periphery and endosomes are enriched, as might have been expected. This validates that the ER-retained mutant could be used to unearth interactions occurring in later steps of vesicular secretion (**Figure 3G**).

Emphasizing on the proteins of interest, we examined which trafficking-related proteins are upregulated, downregulated or not significantly changed between UapA-DYDY and the wild UapA transporter (**Supplementary Table S4 and S5)**. 34 out of the 70 previously annotated trafficking factors of UapA interactome were increased in the wild type transporter relative to UapA-DYDY. These might correspond to interactions that mediate or promote UapA sorting at ERes and/or after ER-exit. Among the other trafficking factors listed in **Supplementary Table S4,** 21 proteins had not significantly change, and 17 proteins were found to be enriched in UapA-DYDY-TurboID).

The interactions „lost‟ in UapA-DYDY included several Golgi proteins (HypB, Syt1a/b, Gst1, and Age1/2), SNARES (SynA, NyvA, Vam7, Syn8, Sec9, and Vti1), post-Golgi proteins (AP-1 and AP-3 components), ER-PM contact proteins (Tcb1-Tcb3), and most endocytic and ESCRT proteins. This further validates that the use of an ER-retained UapA version for detecting post ER-exit interactions. On the other hand, UapA-DYDY seems to conserve, or even increase its interaction with proteins operating at the ER to early-Golgi sorting of cargoes, such as COPII and COPI components, or ArfA. Other proteins showing increased interaction in the ER-retained mutant included early-Golgi SNARES SedV, BetA and Tlg2, Arf-type Arl3 GTPase, but also RabE and RabS, which are known to operate in post-Golgi secretion or late endosomes, respectively (Pinar & Peñalva, 2021). Whether increased biotinylation of early-Golgi proteins or post-Golgi effectors by UapA-DYDY signify functionally relevant interactions, or is just the result of increased concentration of UapA-DYDY in the ER, and thus increased random association with the early-Golgi due to close proximity, remains to be tested. Notably, interactions with COPI and COPII components were enriched in UapA-DYDY relative to UapA, with the only exception being the Sec24 protein (see Heat maps in **Figure 3F**). In other words, Sec24, the protein that mediates cargo recognition during COPII formation, is enriched in the wt UapA. This is an interesting result as it shows that physiologically relevant interactions can be detected when comparing relative cargo mutants, as in our case. On the other hand, it also shows that interactions of UapA-DYDY with other COPII components is *not* specific, but are probably due to close topological proximity within the ER membrane.

Despite the fact that we used derepressed conditions of *de novo* expression of wild type UapA leading to moderate steady state transporter accumulation in the PM, still a fraction of it might well be degraded by constitutive internalization from the PM and sorting to the vacuole. Indeed, the wt UapA interactome obtained included several endocytic, endosomal and vacuolar proteins. To address this issue, we performed PDB assays coupled with proteomics in a genetic background where wild type UapA-TurboID endocytosis is blocked. More specifically, we used a strain containing a null mutant of ArtA (Δ*artA*), the α-arrestin responsible for UapA ubiquitination and endocytosis (Karachaliou et al., 2013). To distinguish between interactions involved in UapA endocytosis and vacuolar degradation from those mainly involved in anterograde trafficking of the transporter, we compared the PDB results of the wild type and the Δ*artA* UapA-TurboID strain. Particularly, 484 protein hits upregulated in the wild type UapA, were considered putative endocytic interactions. (**Supplementary Table S6**).

In the volcano plot blue, colored dots correspond to the protein hits enriched in the wild type UapA (**Figure 4A**). Certain examples of proteins known to participate in vacuolar sorting, such as Nyv1, Hse1, Vps9 etc., are highlighted. Gene enrichment analysis of the dataset of putative endocytic interactions showed that cellular components, such as endosomes and vacuolar membranes, or biological processes, such as vacuolar transport, were enriched (**Figure 4B**). The results verify that the employment of the Δ*artA* background in our experimental set-up can reveal proteins that are physiologically relevant to UapA endocytosis and turnover.

**Supplementary Table S7** shows all cargo trafficking-related proteins identified in the Δ*artA* background. Intriguingly, besides the expected „lost‟ of endocytic and cargo turnover proteins, we also noticed that several SNAREs (Vam7, Syn8, Vti1, Nyv1 and SynA), specific GEFs (Syt1) or GAPs (Gts1 and Age1/2), and proteins involved in post-Golgi secretion (AP-1 or AP-3 subunits), are also lost (i.e., enriched in wild type UapA-TurboID). This result suggests that some of the putative trafficking interactions of wild type UapA might definitely be random and physiologically irrelevant with the biogenesis of the transporter.

### 3.4 Overall validation of UapA interactions

Using the wt UapA proteome results, we pre-selected 70 trafficking-related proteins that seem in close proximity with UapA during its biogenesis. This includes both UapA anterograde trafficking from the ER to the PM and endocytosis and sorting to the vacuoles for degradation. Using the proteome of an ER-retained version of UapA, we showed that 34 out the 70 proteins were in close proximity with UapA after its exit from the ER. Using a genetic background where UapA does not undergo endocytosis and vacuolar degradation we further showed that 30 out of the 70 proteins are in close proximity with UapA after the onset of endocytosis and degradation. The three interactomes permitted partial „filtering‟ of interactions occurring during anterograde trafficking from the ER to the PM, which was the original scope of this work. Based on this, **Table 1** shows 25 proteins that might be true interactors of UapA, during its trafficking from the ER to the PM. Among those, here we decided to validate the possible role of CopA^Cop1^, ArfA^Arf1^, Glo3 and Gcs1 in UapA trafficking. As CopA is an essential component of COPI complex, recruited by ArfA at the interphase of ER with early Golgi, and Glo3 and Gcs1 are known to be regulators of ArfA, our goal was to test whether UapA anterograde trafficking depends on COPI/Arf1.

**Table 1.** Selected proteins interactors of UapA during its trafficking from the ER to the PM. Protein nomenclature of *Saccharomyces cerevisiae* and *A. nidulans* is used.

| ***Aspergillus nidulans***<br>**accession codes in**<br>**FungiDB** | **Systematic/Protein name** | **Trafficking Process** |
| --- | --- | --- |
| AN0411 | Sar1 | COPII coat initiation |
| AN0261 | Sec23 | COPII coat |
| AN3720 | Sec24 | COPII coat |
| AN4317 | Sec13 | COPII coat |
| AN6257 | Sec31 | COPII coat |
| AN6615 | Sec16 | COPII regulation |
| AN11127 | Sec12 | Sar1 GEF |
| AN4724 | Sec1 | SM complex |
| AN3026 | CopA/Cop1 | COPI coat |
| AN1177 | Sec26 | COPI coat |
| AN0922 | Ret2 | COPI coat |
| AN5972 | Sec27 | COPI coat |
| AN5912 | Arl1 | Early/late Golgi sorting |
| AN1126 | ArfA/Arf1 | Early/late Golgi sorting |
| AN9526 | SedV/Sed5 | Tethering (t-SNARE) |
| AN6709 | HypB/Sec7 | Arf1 effector (GEF) |
| AN5127 | BetA/Bet1 | Tethering (t-SNARE) |
| AN2048 | TlgB/Tlg2 | Tethering (t-SNARE) |
| AN4724 | Sec1 | Vesicle docking |
| AN2419 | Sec9 | Tethering (t-SNARE) |
| AN6033 | Glo3 | Arf1 effector (GAP) |
| AN2222 | Gcs1 | Arf1 effector (GAP) |
| AN0347 | RabE/Ypt31/32 | Post-Golgi/ endosome sorting |
| AN0089 | RabS/Ypt7 | Late endosome sorting |
| AN9149 | Tbc1-3 | ER-PM contact sites |

### 3.5 CopA is essential for anterograde cargo trafficking, germination and growth

Among significantly enriched protein interactions of UapA we found several components of the COPI coatomer (CopA^Cop1^, Ret2, Sec26, Sec27). Early studies in yeast have suggested that COPI is primarily involved in retrograde transport of recycling proteins returning from the Golgi to the ER (Pelham, 1994; Letourneur et al., 1994; Cosson and Letourneur, 1997). In contrast to these data, other studies have shown that COPI vesicles carry both anterograde and retrograde cargo (Orci et al., 1997) and can bud from a variety of cellular membranes including the ER, vesicular tubular transport complexes involved in ER to Golgi transport, the Golgi complex and endosomes (Bednarek et al., 1995; Scales et al., 1997; Malhotra et al., 1989; Whitney et al., 1995; Aniento et al., 1996). In a recent article, the ER was shown to produce an interwoven tubular network (identified as ERes) extending microns along microtubules while still connected to the ER by a thin neck, where COPII localizes and dynamically regulates cargo partitioning from the ER (Weigel et al., 2021). In this report, COPI was found to act distally from COPII, escorting the detached, accelerating tubular entity on its way to joining the Golgi apparatus. The distinct functions of COPII and COPI have also been supported by another recent article which showed that only a subpopulation of COPI is localized to ERes, where a COPII collar defines the boundary between ER and ERes exit site and does not coat cargo vesicular carriers (Shomron et al., 2021). Still however, various aspects of cargo ER exit remain unclear (Phuyal and Farhan, 2021).

The possible interaction of COPI with UapA, a cargo that is not known to undergo retrograde trafficking at the ER-Golgi interface, and which seems to exit the ER in specific COPII-dependent vesicles bypassing the Golgi, appeared as interesting to validate. In *A. nidulans,* CopA^Cop1^ has been previously shown to localize and function at the early Golgi (Breakspear et al., 2007; Hernández-González et al., 2019). The same report proposes that the role of COPI is to mediate intra-Golgi retrograde traffic in driving cisternal maturation. Given that CopA is essential for growth and survival, we constructed a controllable knock-down (KD) strain where CopA expression could be tightly repressed using that *thiA* promoter, as previously described for all essential genes of the *A. nidulans* trafficking pathway (Dimou et al., 2020). The *thiA_p_-copA* strain can grow nearly normally in the absence of thiamine from the medium, but cannot germinate and form colonies when thiamine is added for the onset of growth (**Figure 5A**). Relative western blot analysis showed that CopA is indeed tightly repressed in the presence of thiamine (**Figure 5B**).

We crossed the *thiA_p_-copA* strain with a strain expressing UapA tagged with GFP, but also with a strain expressing the v-SNARE SynA-GFP, which is a standard conventional cargo sorted to the PM via the Golgi. Selected progeny containing the KD of *copA* and the GFP-tagged cargoes were used to investigate the role of CopA in membrane cargo trafficking. **Figure 5C** shows the relative epifluorescence analysis (for details of experimental conditions see Materials and methods). Transcriptional repression of CopA, led to dramatic mislocalization of both UapA and SynA from their normal anti-polar and polar PM localization, respectively. More specifically, GFP-SynA labeled mostly the ER, whereas UapA-GFP marked a cytoplasmic haze. This result suggests that CopA is critical for ER-exit and anterograde trafficking of *de novo* made membrane cargoes, as the expression of these cargoes was elicited after having established repression of CopA synthesis. Based on this observation, we then tested whether UapA co-resides with CopA during its sorting to the PM. Colocalization analysis of neosynthesized UapA-GFP with CopA-mRFP (**Figure 5D**) showed that the two signals did not overlap substantially, as verified by the quantitative assessment of the signals‟ correlation (PCC = 0.24 ± 0.06, in n = 11 cells). However, in some cases (**Figure 5D** time series) we noticed a transient signal overlap, in line with the transient nature of cargo ER export, which is very difficult to be captured using widefield epifluorescence microscopy. To further investigate whether the role of CopA on anterograde cargo trafficking is direct, rather than through an effect on COPII biogenesis, we tested the localization of Sec24-GFP in the *thiA_p_-copA* genetic background under thiamine repression conditions (for strain construction see Materials and Methods). **Figure 5E** shows that repressing CopA expression did not affect the formation of Sec24-labeled fluorescent foci. Overall, CopA seems to be essential for the trafficking of both UapA and SynA by operating at a step downstream of COPII formation.

### 3.6 ArfA is essential for anterograde cargo trafficking, germination and growth

Given that COPI is recruited onto membranes by members of the Arf family of small GTPases, we also examined whether the UapA interactome included significantly enriched Arfs. ArfA^Arf1^ and the Arf-like Arl3 were found to be significantly enriched in the interactomes of both wt and ER-retained version of UapA. Arf1 is known to drive COPI-mediated vesicle budding needed for cargo trafficking and secretion, through its role in the Golgi (Jackson and Bouvet, 2014). Arf1 also regulates the formation of clathrin-coated vesicles at the t*rans*-Golgi network by promoting the recruitment of adaptor-protein complexes AP-1, AP-3, AP-4, and the GGAs from the cytosol onto membranes (D’Souza-Schorey and Chavrier, 2006; Sztul et al., 2019). In *A. nidulans*, ArfA has been shown to be an essential protein for germination and growth, via its localization and function in the Golgi network (Lee and Shaw, 2008). Less is known on the role of Arl3 proteins, which are probably related to traffic from endosomes to the Golgi.

To validate the role of ArfA in UapA trafficking we constructed a knock-down (KD) strain where ArfA expression could be tightly repressed, using the *thiA* promoter, as described above for CopA. The *thiA_p_-arfA* strain grows normally in the absence of thiamine from the medium, but does not germinate and cannot form colonies in the presence of thiamine (**Figure 6A** and **6B**). Western blot analysis confirmed that ArfA can be tightly repressed in the presence of thiamine (**Figure 6C**).

We crossed the *thiA_p_-arfA* strain with a strain expressing UapA-GFP or mCH-SynA and studied cargo trafficking in selected progeny, as described for CopA (for strain construction see Materials and Methods). **Figure 6C** shows that transcriptional repression of ArfA, led to mislocalization of both UapA and SynA from their normal PM localization. mCH-SynA and UapA-GFP marked cytoplasmic aggregates/foci of variable size. This result suggests that ArfA, similarly to CopA, is critical for ER-exit and anterograde trafficking of *de novo* made UapA or SynA.

### 3.7 Glo3 or Gcs1 are little critical for growth and cargo trafficking

The regulation of Arfs is governed by GEFs (Guanine Nucleotide Exchange Factors) and GAPs (GTPase Activating Proteins). In their GTP-bound state, which is facilitated by GEFs, Arfs are recruited to membranes. They interact with COPI and influence cargo selection, vesicle scission, and vesicle uncoating. GTP hydrolysis on Arfs is necessary for the dissociation of coat proteins from transport vesicles and is mediated by GAPs. GTP hydrolysis on Arf1 is also required for cargo packaging, and Arf GAPs might function to couple cargo sorting with vesicle formation (Jackson and Bouvet, 2014).

In the interactome of UapA, two highly enriched ArfA GAP proteins were identified: Glo3 (with a difference of >4 compared to the control strain) and Gcs1 (with a difference of >7). Consequently, we investigated the role of both proteins in UapA trafficking by constructing and analyzing the respective null mutants (Δ*glo3* and Δ*gcs1*) as described in the Materials and Methods section. Interestingly, the double mutant Δglo3 Δgcs1 is lethal, as evidenced by heterokaryon rescue (not shown), in line with reports in yeast cells (Poon et al., 1999). **Figure 7A** illustrates the growth phenotypes of Δ*glo3* and Δ*gcs1* on various media. Δ*glo3* exhibits a moderate modification in conidiation and colony growth at 37°C but not at 25°C, whereas Δ*gcs1* does not display any apparent mutant phenotype.

**Figure 7.**
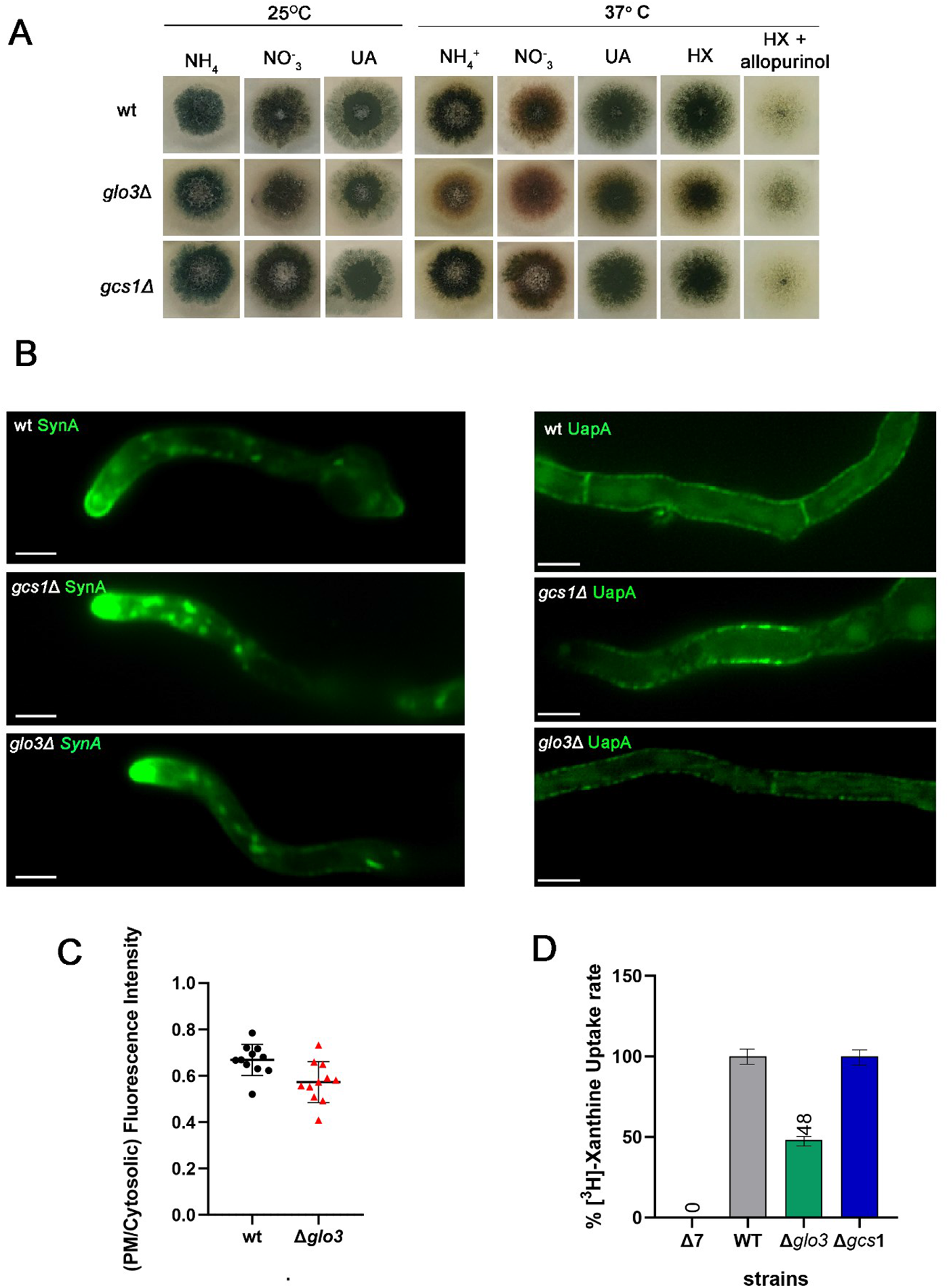
Glo3 or Gcs1 are little critical for growth and cargo trafficking. **A.** Growth tests of knock-out strains Δ*glo3* and Δ*gcs1* on selected N sources at 25°C and 37°C compared to a wild-type isogenic control strain (wt). N sources shown are: 10 mM ammonium tartrate (NH_4_^+^), 10 mM sodium nitrate (NO_3_^-^), 0.5 mM uric acid (UA), 0.5 mM hypoxanthine (HX) and 3 μM allopurinol. Deletion of *gcs1* has no phenotypic effect on all conditions tested, while Δ*glo3* shows a minor growth defect at 37°C. In the presence of the cytotoxic analogue, allopurinol, the *glo3* null mutant seems to have better sporulation relative to the wt strain, indicating reduced function of UapA. **B.** Epifluorescence microscopy analysis of the subcellular localization of UapA and SynA in genetic backgrounds where *glo3* or *gcs1* are deleted, at 25°C. In Δ*glo3* and Δ*gcs1* mutants, both cargoes reach the PM. However in *Δglo3,* although UapA reaches the PM, the fluorescence of the transporter to the PM seems to be lower **C.** Scatter plot depicting the UapA-GFP PM/cytosolic intensity ratios in a wt and a Δ*glo3* genetic background. Mean PM/cytosolic intensity ratios are 0.67 ± 0.02 and 0.57 ± 0.03, respectively. To test the significance of differences, an unpaired t-test was performed, which verified the significant difference (P = 0.0097). Biological/technical replicates: 2/11 for each condition. **D.** Relative ^3^H-xanthine (0.3 μM) transport rates of the null mutants, Δ*glo3* and Δ*gcs1*, analyzed and expressed as percentages of initial uptake rates (V) compared to the wild type UapA rate. Δ7 is a strain lacking all 7 major nucleobase transporters, consisting of the negative control for the transport assays. The time point used for the initial uptake rate is 1 min. Results are averages of 3 measurements using the same concentration of spores for each strain. SD was less than 20%.

To quantify the possible effect of these mutants on UapA localization, we also conducted growth test assays in the presence of the cytotoxic analogue allopurinol. Allopurinol is taken up by the UapA transporter and drastically inhibits xanthine dehydrogenase, hxA, an enzyme necessary for the catabolism of hypoxanthine (HX) (Elion, 1989; Pantazopoulou & Diallinas, 2007). Therefore, when allopurinol is present, HX cannot be utilized as the sole nitrogen source. This is a test more sensitive for the measuring the activity of UapA compared to growth on its physiological substrates as N sources. In **Figure 7A**, the *glo3* null mutant shows slightly improved sporulation, relative to the wild type, in media containing HX and allopurinol, indicating reduced uptake of the cytotoxic compound. This result suggests that Δ*glo3* may moderately affect the quantity of UapA transporter on the plasma membrane.

To test directly the effects of Glo3 and Gcs1 on cargo trafficking, we crossed both null mutants with a strain expressing UapA-GFP or GFP-SynA, and analyzed the resulting progeny. **Figure 7B** demonstrates that the PM localization of both cargoes is unaffected at 25°C, except from an apparent reduction of the fluorescent intensity of UapA-GFP at the PM. This moderate reduction was further confirmed by fluorescence quantification (**Figure 7C**). To validate the reduced localization of UapA-GFP to the PM, we also performed transport assays with radiolabeled xanthine, the substrate of UapA. **Figure 7D** shows that the capacity of UapA transport is reduced by half in Δ*glo3*, but remains nearly normal in Δ*gcs1*, consistent with the growth tests and epifluorescence microscopy. Overall, our results suggest a physiologically relevant interaction between UapA and Glo3, but not with Gcs1.

In *A. nidulans,* ArfA is also regulated by GEFs GeaA^Gea1^ and HypB^Sec7^ at the early-Golgi or late-Golgi, respectively. The interactomes of UapA did not include GeaA, but there was significant enrichment of HypB in the interactome of wild type UapA. We have previously provided strong evidence that none of these GEFs is needed for proper UapA trafficking to the PM, but they are both essential for the trafficking of SynA and other Golgi-dependent cargoes (Dimou et al., 2020). We thus concluded that enrichment of HypB in the UapA interactome is very probably a physiologically irrelevant interaction,

## 4. Conclusions

The mechanism of anterograde trafficking of eukaryotic polytopic transmembrane proteins, such as transporters, channels or receptors, which constitute the most abundant types of PM cargoes, has not been studied in a systematic and rigorous manner. Standard methods for identifying protein-protein interaction, as two-hybrid approaches, affinity capture, co-fractionation, FRET or bifluorescence, failed to identify trafficking proteins interacting physically and thus mediating the translocation of membrane cargoes from the ER to the PM. In the *S. cerevisiae* database (https://www.yeastgenome.org/), for example, we could hardly find any transporter interacting physically with Sec24, the main cargo receptor of newly made membrane proteins at the ER. This is probably due to the transient and super-rapid nature of these interactions, estimated to be at the millisecond level.

In the present work we employed PDB assays coupled with proteomics to identify proteins interacting with the UapA transporter, a previously reported fast-moving cargo that seemingly bypasses the Golgi on its way from the ER to the PM (Dimou et al., 2020; 2022). Combining results from three different proteomes and under the light of current knowledge on proteins essential (or redundant) for UapA trafficking, we came to a list of 25 proteins that might be proximal to UapA in the course of its trafficking to the PM. Some of them, such as COPII components, are expected to be in close proximity with UapA, given our previous studies that showed direct dependence of UapA localization to the PM on functional Sec24 and Sec13 proteins (Dimou et al., 2020; 2022). However, other proximity interactions identified herein included several early-Golgi proteins (e.g., CopA, ArfA, Glo3, Gst1, SedV, BetA, etc.), and a few late-Golgi (HypB^Sec7^) or post-Golgi (RabE) proteins, contrasting the idea that UapA completely bypasses Golgi functioning.

The identification of specific early-Golgi proteins in our PDB assays signify that UapA localization might indeed be partially dependent on sorting to a very early ER-Golgi intermediate compartment (ERGIC). Alternatively, several of these interactions might be physiologically irrelevant and occur simply due to very close proximity of ERes to early-Golgi structures. Similarly, other unexpected UapA interactions, due to previous genetic evidence (Dimou et al., 2020; 2022), as for example with RabE, might signify either a new function of RabE into Golgi-independent cargo trafficking, or simply be an artifact of the technique. Notably, however, several key Golgi-related proteins (e.g., RabO) were not among those found to be proximal to UapA, which is in line with the concept of Golgi-bypass of UapA (Dimou et al., 2020).

Here, we decided to validate the functional role of CopA and ArfA in UapA trafficking, as these two factors regulate cargo secretion through the early-Golgi and recent reports supported the direct involvement of COPI is anterograde trafficking of membrane cargoes (Shomron et al., 2021; Weigel et al., 2021). We provided evidence that COPI (i.e., CopA) is indeed essential for trafficking of both cargoes tested (UapA and SynA) via a mechanism operating downstream of COPII. As retrograde and anterograde pathways are COPI-dependent and tightly coupled, we cannot exclude that the requirement of COPI in cargo trafficking could be due to a block in retrograde sorting of UapA or SynA to the ER. In line with the essentiality of COPI in UapA and SynA trafficking, experimentally supported herein, we also showed that ArfA is also essential for localization of these cargoes to the PM. Notably, however, the two known GAPs expected to regulate ArfA in *A. nidulans* were shown here to have no (Gcs1) or minor (Glo3) effect on UapA or SynA sorting to the PM. This suggests that ArfA might be crucial for specific cargo sorting via distinct interactions with GAPs or GEFs.

The current work identified several new candidates as possible UapA trafficking effectors operating after ER-exit. Among those, Sec1, Sec9 and Tbc1/3 are particularly attractive, as their function is well compatible with direct cargo sorting from an early-secretory compartment to the PM. Sec1 is an SM-like protein involved in docking and fusion of exocytic vesicles (Südhof and Rothman 2009; Morgera et al., 2011) via binding to assembled PM SNARE complexes. Sec9 is a PM t-SNARE protein required for secretory vesicle fusion (Geert van den Bogaart and Jahn, 2011; Dubuke et al., 2015). Finally, Tcb1/3 is lipid-binding ER tricalbin involved in cortical ER-plasma membrane tethering (Manford et al., 2012; Collado et al., 2019).

In conclusion, we are aware that the present PDB system suffers from lack of functional specificity, as many interactions identified are simply irrelevant to UapA trafficking. In line with this, the entire UapA interactome included approximately 34% of metabolite interconversion enzymes (**Figure 2B**), unrelated to membrane cargo trafficking. Additionally, we identified several interactions with cytosolic proteins involved in post-Golgi secretion, such as AP-1 and AP-3 components, which have been rigorously shown to be fully dispensable for UapA trafficking (Martzoukou et al., 2018; Dimou et al., 2020). To increase the specificity of our system, we envision performing PDB assays with proteomes obtained from mutants blocked in distinct steps of Golgi or post-Golgi functioning, thus being able to track the sequential steps of distinct cargoes that traffic either via the conventional Golgi-dependent route or via Golgi-bypass. Overall, acknowledging the advantages, the limitations and the future prospects of the current methodology, we conclude that PDB might be a useful tool for unraveling transient interactions in filamentous fungi.

## Supporting information

Supplementary Georgiou et al.

## Abbreviations

UapA-DYDY-TurboID: UapA-Asp-Tyr-Asp-Tyr/AAAA-TurboID
ER: endoplasmic reticulum
PDB: proximity dependent biotinylation
PM: plasma membrane
Wt: wild type
GAP: GTPase activating protein
GEF: guanine nucleotide exchange factor
N: nitrogen
Thi: thiamine
LC-MS/MS: liquid chromatography coupled with tandem mass spectrometry.

## Data availability

All strains and primers used in this study are listed in **Appendix Tables S1, S2**. The mass spectrometry proteomics data are available in the ProteomeXchange Consortium via the PRIDE partner repository with the following dataset identifiers:

1. PXD043934 **Username:** Password: 9SnPcHqV

2. PXD043941 **Username:** Password: 0gwq3Smi

3. PXD043944 **Username:** Password: i08neOut

## Supplementary Material

Provided separately as excel files

## Author contributions

X.G: investigation, visualization, data curation, methodology, validation, formal analysis, software, conceptualization, writing-review/editing, S.D: investigation, validation, writing-review/editing, formal analysis, visualization, M.S: conceptualization, investigation G.D: conceptualization, supervision, writing-original draft, project administration, funding acquisition.

## Acknowledgments

We are grateful of Dr. Sotirs Amilis for constructs and strains concerning CopA and ArfA. We also thank Prof. Uygar Tazebay for the TurboID clones and his ideas on PDB assays, and Prof. Panagiotis Moschou on his suggestions on PDB assays. This work was supported by an HFRI research grant (KE 18458), the Fondation Santé. We acknowledge support of this work by the project “The Greek Research Infrastructure for Personalised Medicine (pMedGR)” (MIS 5002802) which is implemented under the Action “Reinforcement of the Research and Innovation Infrastructure”, funded by the Operational Programme “Competitiveness, Entrepreneurship and Innovation” (NSRF 2014-2020) and co-financed by Greece and the European Union (European Regional Development Fund).

## Appendix

**Table S1.** *A. nidulans* strains used in this study. All strains carry the *veA1* mutation affecting sporulation. *pabaA1*, *pyroA4*, *riboB2*, *argB2* and *pyrG89* are auxotrophic mutations for p-aminobenzoic acid, pyridoxine, riboflavin, arginine, and uracil/uridine, respectively.

| Strain | Genotype | Source |
| --- | --- | --- |
| wt | *pabaA1* | wild-type reference strain |
| TNO2A7 | $\Delta nkuA::argB$ *pyrG89 pyroA4 riboB2* | Nayak et al., 2006 |
| $\Delta 3$ | $\Delta uapA \Delta uapC::AFpyrG \Delta azgA pBS-argB pabaA1 argB2$ | Pantazopoulou et al., 2007 |
| $\Delta 7$ | $\Delta uapA \Delta uapC::AFpyrG \Delta azgA \Delta fcyB::argB \Delta furD::AFriboB \Delta furA::AFriboB \Delta cntA::AFriboB pantoB100 pabaA1$ | Kryptou and Diallinas, 2014 |
| UapA-TurboID | $(pan510exp)-uapA-Turbo \Delta uapA \Delta uapC::AFpyrG \Delta azgA pabaA1 argB2$ | This study |
| UapA-DYDY-TurboID | $(pan510exp)-uapA-D44A/Y45A/D46A/Y47A-Turbo \Delta uapA \Delta uapC::AFpyrG \Delta azgA pabaA1 argB2$ | This study |
| UapA-TurboID $\Delta artA$ | $(pan510exp)-uapA-Turbo \Delta uapA \Delta uapC::AFpyrG \Delta azgA \Delta artA::AFriboB pabaA1$ | This study |
| thiAp-copA | *thiA<sub>p</sub>-copA(AN3026)::AFpyrG $\Delta nkuA::argB$ *pyrG89 riboB2 pyroA4** | This study |
| thiAp-<sup>FLAG</sup>copA | *thiA<sub>p</sub>-<sup>FLAG</sup>copA(AN3026)::AFpyrG $\Delta nkuA::argB$ *pyrG89 riboB2 pyroA4** | This study |
| UapA-GFP thiAp-copA | $\Delta uapA::uapA-GFP^L thiA_p-copA(AN3026)::AFpyrG \Delta nkuA::argB pabaA1$ | This study |
| GFP-SynA thiAp-copA | $alcA_p-GFP-synA::AFpyrG thiA_p-copA(AN3026)::AFpyrG \Delta nkuA::argB/bar pyroA4$ | This study |
| UapA-GFP CopA-mRFP | $\Delta uapA::alcA_p-uapA-GFP^L::AFriboB copA(AN3026)-(5xGA)mRFP::AFpyrG \Delta nkuA::argB pyrG89 pyroA4 riboB2$ | This study |
| Sec24-GFP thiAp-copA | $sec24-(5xGA)GFP::AFpyrG \Delta nkuA::argB pyrG89 thiA_p-copA(AN3026)::AFpyrG pyroA4$ | This study |
| thiAp-arfA-GFP | $AFriboB::thiA_p-arfA(AN1126)-(5xGA)GFP::AFpyrG \Delta nkuA::argB pyrG89 pyroA4 riboB2$ | This study |
| UapA-GFP thiAp-arfA | $thiA_p-arfA(AN1126)::AFpyrG \Delta uapA::uapA-GFP^L \Delta nkuA::argB pyrG89 pyroA4$ | This study |
| mCh-SynA thiAp-arfA | $thiA^P-arfA(AN1126)::AFpyrG alcA_p-mCH-synA::AFpyroA4 \Delta nkuA::argB pyrG89 pyroA4$ | This study |
| UapA-GFP $\Delta glo3$ | $\Delta glo3(AN6033)::AFpyrG \Delta uapA::uapA-GFP^L::AFriboB \Delta nkuA::argB pyroA4 riboB2 pyrG89$ | This study |
| UapA-GFP $\Delta gcs1$ | $\Delta gcs1(AN2222)::AFpyrG \Delta uapA::uapA-GFP^L::AFriboB \Delta nkuA::argB pyroA4 riboB2 pyrG89$ | This study |
| GFP-SynA $\Delta glo3$ | $\Delta glo3(AN6033)::AFpyrG alcA_p-GFP-synA::AFpyrG \Delta nkuA::argB pyroA4 riboB2 pyrG89$ | This study |
| GFP-SynA $\Delta gcs1$ | $\Delta gcs1(AN2222)::AFpyrG alcA_p-GFP-synA::AFpyrG \Delta nkuA::argB pyroA4 riboB2 pyrG89$ | This study |

**Table S2.**
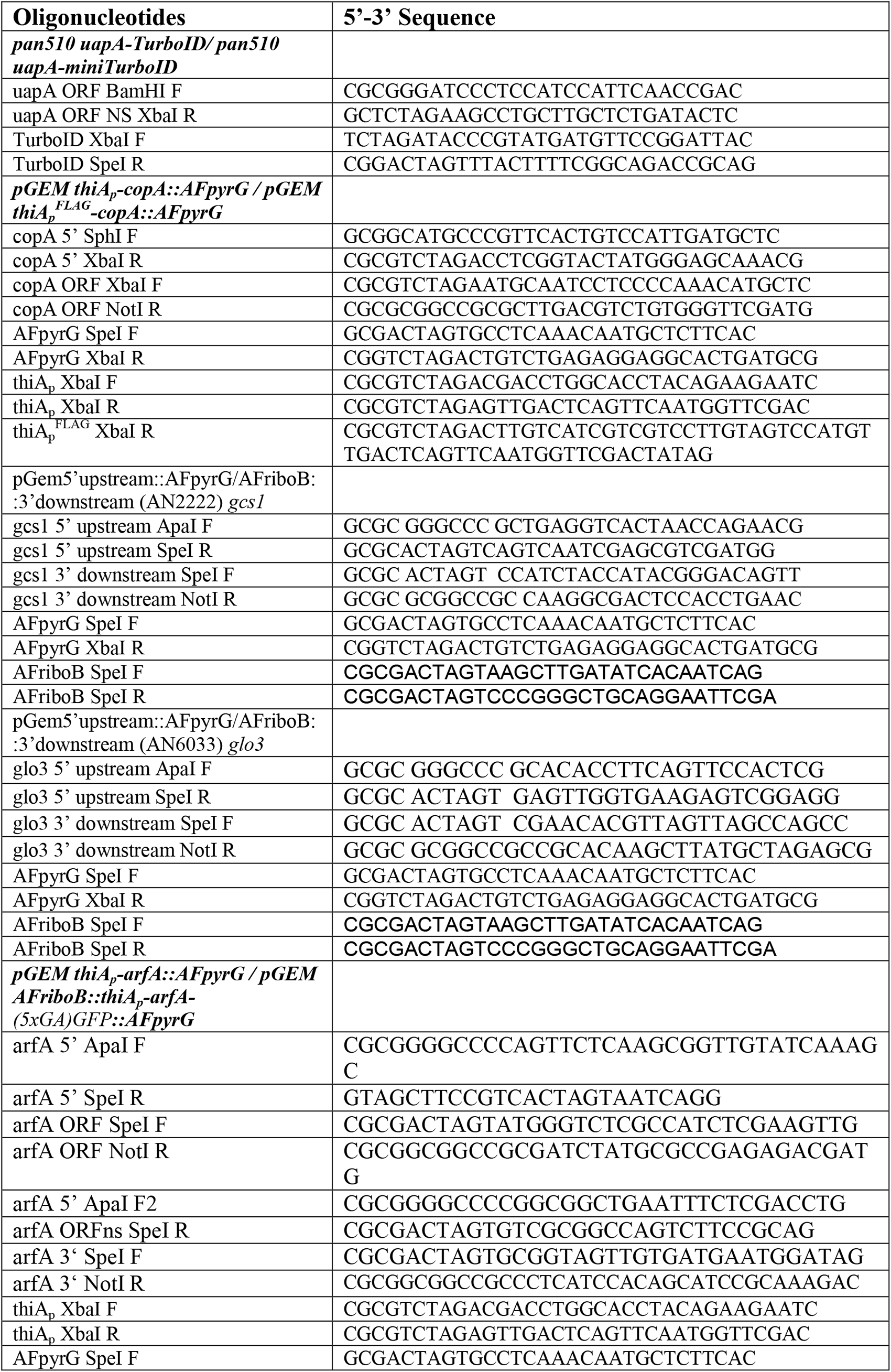
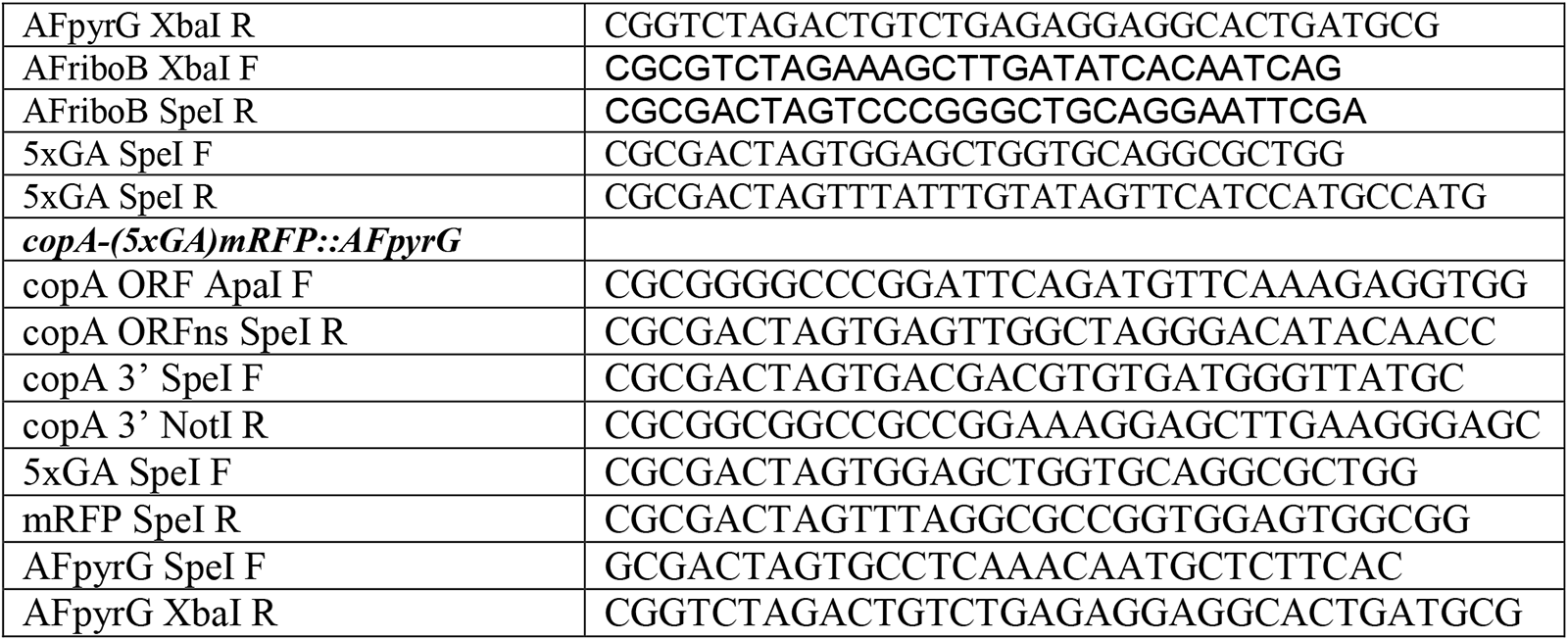
Oligonucleotides used in this study for cloning purposes.

## References

Alguel Y, Amillis S, Leung J, Lambrinidis G, Capaldi S, Scull NJ, Craven G, Iwata S, Armstrong A, Mikros E, Diallinas G, Cameron AD, Byrne B. Structure of eukaryotic purine/H(+) symporter UapA suggests a role for homodimerization in transport activity. Nat Commun. 2016 Apr 18;7:11336. doi: 10.1038/ncomms11336.

Aniento F, Gu F, Parton RG, Gruenberg J. An endosomal beta COP is involved in the pH-dependent formation of transport vesicles destined for late endosomes. J Cell Biol. 1996 Apr;133(1):29–41. doi: 10.1083/jcb.133.1.29.

Apostolaki, A., Harispe, L., Calcagno-Pizarelli, A.M., Vangelatos, I., Sophianopoulou, V., Arst Jr, H.N., Peñalva, M.A., Amillis, S. and Scazzocchio, C. (2012), *Aspergillus nidulans* CkiA is an essential casein kinase I required for delivery of amino acid transporters to the plasma membrane. Molecular Microbiology, 84: 530–549. https://doi.org/10.1111/j.1365-2958.2012.08042.x

Arora D, Abel NB, Liu C, Van Damme P, Yperman K, Eeckhout D, Vu LD, Wang J, Tornkvist A, Impens F, Korbei B, Van Leene J, Goossens A, De Jaeger G, Ott T, Moschou PN, Van Damme D. Establishment of Proximity-Dependent Biotinylation Approaches in Different Plant Model Systems. Plant Cell. 2020 Nov;32(11):3388–3407. doi: 10.1105/tpc.20.00235.

Basenko EY, Pulman JA, Shanmugasundram A, Harb OS, Crouch K, Starns D, Warrenfeltz S, Aurrecoechea C, Stoeckert CJ Jr, Kissinger JC, Roos DS, Hertz-Fowler C. FungiDB: An Integrated Bioinformatic Resource for Fungi and Oomycetes. J Fungi (Basel). 2018 Mar 20;4(1):39. doi: 10.3390/jof4010039.

Bednarek SY, Ravazzola M, Hosobuchi M, Amherdt M, Perrelet A, Schekman R, Orci L. COPI- and COPII-coated vesicles bud directly from the endoplasmic reticulum in yeast. Cell. 1995 Dec 29;83(7):1183–96. doi: 10.1016/0092-8674(95)90144-2

Bonifacino JS, Glick BS. The mechanisms of vesicle budding and fusion. Cell. 2004 Jan 23;116(2):153–66. doi: 10.1016/s0092-8674(03)01079-1.

Branon TC, Bosch JA, Sanchez AD, Udeshi ND, Svinkina T, Carr SA, Feldman JL, Perrimon N, Ting AY. Efficient proximity labeling in living cells and organisms with TurboID. Nat Biotechnol. 2018 Oct;36(9):880–887. doi: 10.1038/nbt.4201.

Breakspear A, Langford KJ, Momany M, Assinder SJ. CopA:GFP localizes to putative Golgi equivalents in Aspergillus nidulans. FEMS Microbiol Lett. 2007 Dec;277(1):90–7. doi: 10.1111/j.1574-6968.2007.00945.x.

Camacho, C., Coulouris, G., Avagyan, V., Ma, N., Papadopoulos, J., Bealer, K., & Madden, T. L. (2009). BLAST+: Architecture and applications. BMC Bioinformatics, 10(1). https://doi.org/10.1186/1471-2105-10-421.

Catherine L. Jackson, Samuel Bouvet; Arfs at a Glance. J Cell Sci 1 October 2014; 127 (19): 4103–4109. doi: https://doi.org/10.1242/jcs.144899.

Cherry JM, Hong EL, Amundsen C, Balakrishnan R, Binkley G, Chan ET, Christie KR, Costanzo MC, Dwight SS, Engel SR, Fisk DG, Hirschman JE, Hitz BC, Karra K, Krieger CJ, Miyasato SR, Nash RS, Park J, Skrzypek MS, Simison M, Weng S, Wong ED (2012) Saccharomyces Genome Database: the genomics resource of budding yeast. Nucleic Acids Res. Jan;40(Database issue):D700-5. [PMID: 22110037]

Cho KF, Branon TC, Udeshi ND, Myers SA, Carr SA, Ting AY. Proximity labeling in mammalian cells with TurboID and split-TurboID. Nat Protoc. 2020 Dec;15(12):3971–3999. doi: 10.1038/s41596-020-0399-0.

Cohen N, Aviram N, Schuldiner M. A systematic proximity ligation approach to studying protein-substrate specificity identifies the substrate spectrum of the Ssh1 translocon. EMBO J. 2023 Jun 1;42(11):e113385. doi: 10.15252/embj.2022113385.

Collado J, Kalemanov M, Campelo F, Bourgoint C, Thomas F, Loewith R, Martínez-Sánchez A, Baumeister W, Stefan CJ, Fernández-Busnadiego R. Tricalbin-Mediated Contact Sites Control ER Curvature to Maintain Plasma Membrane Integrity. Dev Cell. 2019 Nov 18;51(4):476–487.e7. doi: 10.1016/j.devcel.2019.10.018.

Cosson P, Letourneur F. Coatomer (COPI)-coated vesicles: role in intracellular transport and protein sorting. Curr Opin Cell Biol. 1997 Aug;9(4):484–7. doi: 10.1016/s0955-0674(97)80023-3.

Diallinas G, Martzoukou O. Transporter membrane traffic and function: lessons from a mould. FEBS J. 2019 Dec;286(24):4861–4875. doi: 10.1111/febs.15078.

Diallinas G. Dissection of Transporter Function: From Genetics to Structure. Trends Genet. 2016 Sep;32(9):576–590. doi: 10.1016/j.tig.2016.06.003.

Dimou S, Dionysopoulou M, Sagia GM, Diallinas G. Golgi-Bypass Is a Major Unconventional Route for Translocation to the Plasma Membrane of Non-Apical Membrane Cargoes in Aspergillus nidulans Front Cell Dev Biol. 2022 Apr 7;10:852028. doi: 10.3389/fcell.2022.852028.

Dimou S, Georgiou X, Sarantidi E, Diallinas G, Anagnostopoulos AK. Profile of Membrane Cargo Trafficking Proteins and Transporters Expressed under N Source Derepressing Conditions in Aspergillus nidulans. J Fungi (Basel). 2021 Jul 14;7(7):560. doi: 10.3390/jof7070560.

Dimou S, Martzoukou O, Dionysopoulou M, Bouris V, Amillis S, Diallinas G. Translocation of nutrient transporters to cell membrane via Golgi bypass in Aspergillus nidulans. EMBO Rep. 2020 Jul 3;21(7):e49929. doi: 10.15252/embr.201949929.

D’Souza-Schorey C, Chavrier P. ARF proteins: roles in membrane traffic and beyond. Nat Rev Mol Cell Biol. 2006 May;7(5):347–58. doi: 10.1038/nrm1910

Dubuke ML, Maniatis S, Shaffer SA, Munson M. The Exocyst Subunit Sec6 Interacts with Assembled Exocytic SNARE Complexes. J Biol Chem. 2015 Nov 20;290(47):28245–28256. doi: 10.1074/jbc.M115.673806.

Dyballa N, Metzger S. Fast and sensitive colloidal coomassie G-250 staining for proteins in polyacrylamide gels. J Vis Exp. 2009 Aug 3;(30):1431. doi: 10.3791/1431.

Elion GB (1989) The purine path to chemotherapy. Science 244: 41–47.

Emr S, Glick BS, Linstedt AD, Lippincott-Schwartz J, Luini A, Malhotra V, Marsh BJ, Nakano A, Pfeffer SR, Rabouille C, Rothman JE, Warren G, Wieland FT. Journeys through the Golgi--taking stock in a new era. J Cell Biol. 2009 Nov 16;187(4):449–53. doi: 10.1083/jcb.200909011.

Fenech EJ, Cohen N, Kupervaser M, Gazi Z, Schuldiner M. A toolbox for systematic discovery of stable and transient protein interactors in baker’s yeast. Mol Syst Biol. 2023 Feb 10;19(2):e11084. doi: 10.15252/msb.202211084.

Geert van den Bogaart, Reinhard Jahn, Counting the SNAREs needed for membrane fusion, *Journal of Molecular Cell Biology*, Volume 3, Issue 4, August 2011, Pages 204–205, https://doi.org/10.1093/jmcb/mjr004

Gillingham AK, Sinka R, Torres IL, Lilley KS, Munro S. Toward a comprehensive map of the effectors of rab GTPases. Dev Cell. 2014 Nov 10;31(3):358–373. doi: 10.1016/j.devcel.2014.10.007.

Gingras AC, Abe KT, Raught B. Getting to know the neighborhood: using proximity-dependent biotinylation to characterize protein complexes and map organelles. Curr Opin Chem Biol. 2019 Feb;48:44–54. doi: 10.1016/j.cbpa.2018.10.017.

Goedhart, J., and Luijsterburg, M.S. (2020). VolcaNoseR is a web app for creating, exploring, labeling and sharing volcano plots. Sci. Rep. 10, 20560. 10.1038/s41598-020-76603-3.

Gournas C, Amillis S, Vlanti A, Diallinas G. Transport-dependent endocytosis and turnover of a uric acid-xanthine permease. Mol Microbiol. 2010 Jan;75(1):246–60. doi: 10.1111/j.1365-2958.2009.06997.x.20002879.

Hernández-González M, Bravo-Plaza I, de Los Ríos V, Pinar M, Pantazopoulou A, Peñalva MA. COPI localizes to the early Golgi in Aspergillus nidulans. Fungal Genet Biol. 2019 Feb;123:78–86. doi: 10.1016/j.fgb.2018.12.003.

Hollstein LS, Schmitt K, Valerius O, Stahlhut G, Pöggeler S. Establishment of in vivo proximity labeling with biotin using TurboID in the filamentous fungus Sordaria macrospora. Sci Rep. 2022 Oct 22;12(1):17727. doi: 10.1038/s41598-022-22545-x.

J. D. Hunter, “Matplotlib: A 2D Graphics Environment”, Computing in Science & Engineering, vol. 9, no. 3, pp. 90–95, 2007

Jahn R, Südhof TC. Membrane fusion and exocytosis. Annu Rev Biochem. 1999;68:863–911. doi: 10.1146/annurev.biochem.68.1.863.

Karachaliou M, Amillis S, Evangelinos M, Kokotos AC, Yalelis V, Diallinas G. The arrestin-like protein ArtA is essential for ubiquitination and endocytosis of the UapA transporter in response to both broad-range and specific signals. Mol Microbiol. 2013 Apr;88(2):301–17. doi: 10.1111/mmi.12184.

Kershberg L, Banerjee A, Kaeser PS. Protein composition of axonal dopamine release sites in the striatum. Elife. 2022 Dec 29;11:e83018. doi: 10.7554/eLife.83018.

Koukaki M, Giannoutsou E, Karagouni A, Diallinas G. A novel improved method for Aspergillus nidulans transformation. J Microbiol Methods. 2003 Dec;55(3):687–95. doi: 10.1016/s0167-7012(03)00208-2.

Krypotou E, Diallinas G. Transport assays in filamentous fungi: kinetic characterization of the UapC purine transporter of Aspergillus nidulans. Fungal Genet Biol. 2014 Feb;63:1–8. doi: 10.1016/j.fgb.2013.12.004.

Lee SC, Shaw BD. Localization and function of ADP ribosylation factor A in Aspergillus nidulans. FEMS Microbiol Lett. 2008 Jun;283(2):216–22. doi: 10.1111/j.1574-6968.2008.01174.x.

Letourneur F, Gaynor EC, Hennecke S, Démollière C, Duden R, Emr SD, Riezman H, Cosson P. Coatomer is essential for retrieval of dilysine-tagged proteins to the endoplasmic reticulum. Cell. 1994 Dec 30;79(7):1199–207. doi: 10.1016/0092-8674(94)90011-6.

Mair A, Xu SL, Branon TC, Ting AY, Bergmann DC. Proximity labeling of protein complexes and cell-type-specific organellar proteomes in *Arabidopsis* enabled by TurboID. Elife. 2019 Sep 19;8:e47864. doi: 10.7554/eLife.47864.

Malhotra V, Serafini T, Orci L, Shepherd JC, Rothman JE. Purification of a novel class of coated vesicles mediating biosynthetic protein transport through the Golgi stack. Cell. 1989 Jul 28;58(2):329–36. doi: 10.1016/0092-8674(89)90847-7.

Manford AG, Stefan CJ, Yuan HL, Macgurn JA, Emr SD. ER-to-plasma membrane tethering proteins regulate cell signaling and ER morphology. Dev Cell. 2012 Dec11;23(6):1129-40. doi: 10.1016/j.devcel.2012.11.004

Marc Larochelle, Danny Bergeron, Bruno Arcand, François Bachand; Proximity-dependent biotinylation mediated by TurboID to identify protein–protein interaction networks in yeast. J Cell Sci 1 June 2019; 132 (11): jcs232249. doi: https://doi.org/10.1242/jcs.232249

Martzoukou O, Diallinas G, Amillis S. Secretory Vesicle Polar Sorting, Endosome Recycling and Cytoskeleton Organization Require the AP-1 Complex in Aspergillus nidulans. Genetics. 2018 Aug;209(4):1121–1138. doi: 10.1534/genetics.118.301240.

Martzoukou O, Karachaliou M, Yalelis V, Leung J, Byrne B, Amillis S, Diallinas G. Oligomerization of the UapA Purine Transporter Is Critical for ER-Exit, Plasma Membrane Localization and Turnover. J Mol Biol. 2015 Aug 14;427(16):2679–96. doi: 10.1016/j.jmb.2015.05.021.

Miguel Hernandez-Gonzalez (2018), Traffic through the Golgi apparatus and apical extension in Aspergillus nidulans, Complutense University of Madrid.

Morgera F, Sallah MR, Dubuke ML, Gandhi P, Brewer DN, Carr CM, Munson M. Regulation of exocytosis by the exocyst subunit Sec6 and the SM protein Sec1. Mol Biol Cell. 2012 Jan;23(2):337–46. doi: 10.1091/mbc.E11-08-0670.

Orci L, Stamnes M, Ravazzola M, Amherdt M, Perrelet A, Söllner TH, Rothman JE. Bidirectional transport by distinct populations of COPI-coated vesicles. Cell. 1997 Jul 25;90(2):335–49. doi: 10.1016/s0092-8674(00)80341-4.

Pantazopoulou A, Glick BS. A Kinetic View of Membrane Traffic Pathways Can Transcend the Classical View of Golgi Compartments. Front Cell Dev Biol. 2019 Aug 6;7:153. doi: 10.3389/fcell.2019.00153.

Pantazopoulou A, Lemuh ND, Hatzinikolaou DG, Drevet C, Cecchetto G, Scazzocchio C, Diallinas G. Differential physiological and developmental expression of the UapA and AzgA purine transporters in Aspergillus nidulans. Fungal Genet Biol. 2007 Jul;44(7):627–40. doi: 10.1016/j.fgb.2006.10.003

Pantazopoulou A, Pinar M, Xiang X, Peñalva MA. Maturation of late Golgi cisternae into RabE(RAB11) exocytic post-Golgi carriers visualized in vivo. Mol Biol Cell. 2014 Aug 15;25(16):2428–43. doi: 10.1091/mbc.E14-02-0710.

Pantazopoulou A. The Golgi apparatus: insights from filamentous fungi. Mycologia. 2016 May-Jun;108(3):603–22. doi: 10.3852/15-309

Pantazopoulou, A., & Diallinas, G. (2007). Fungal Nucleobase Transporters. FEMS Microbiology Reviews, 31(6), 657–675. https://doi.org/10.1111/j.1574-6976.2007.00083.x.

Pelham HR. About turn for the COPs? Cell. 1994 Dec 30;79(7):1125–7. doi: 10.1016/0092-8674(94)90002-7. P

Perez-Riverol, Y., Bai, J., Bandla, C., García-Seisdedos, D., Hewapathirana, S., Kamatchinathan, S., Kundu, D.J., Prakash, A., Frericks-Zipper, A., Eisenacher, M., et al. (2022). The PRIDE database resources in 2022: a hub for mass spectrometry-based proteomics evidences. Nucleic Acids Res. 50, D543–D552. 10.1093/nar/gkab1038.

Phuyal S, Farhan H. Want to leave the ER? We offer vesicles, tubules, and tunnels. J Cell Biol. 2021 June 7;220(6):e202104062. doi: 10.1083/jcb.202104062.

Pinar M, Peñalva MA. The fungal RABOME: RAB GTPases acting in the endocytic and exocytic pathways of Aspergillus nidulans (with excursions to other filamentous fungi). Mol Microbiol. 2021 Jul;116(1):53–70. doi:10.1111/mmi.1471

Poon, P. P. (1999). Retrograde transport from the yeast golgi is mediated by two arf gap proteins with overlapping function. The EMBO Journal, 18(3), 555–564. https://doi.org/10.1093/emboj/18.3.555

Robinson, M. S. (2015). Forty Years of Clathrin-coated Vesicles. In Traffic (Vol. 16, Issue 12). https://doi.org/10.1111/tra.12335.

Rojas, A. M., Fuentes, G., Rausell, A., & Valencia, A. (2012). The Ras protein superfamily: Evolutionary tree and role of conserved amino acids. In Journal of Cell Biology (Vol. 196, Issue 2). https://doi.org/10.1083/jcb.201103008

Rothman, J. Mechanisms of intracellular protein transport. Nature 372, 55–63 (1994). https://doi.org/10.1038/372055a0

Sander H, Wallace S, Plouse R, Tiwari S, Gomes AV. Ponceau S waste: Ponceau S staining for total protein normalization. Anal Biochem. 2019 Jun 15;575:44–53. doi: 10.1016/j.ab.2019.03.010.

Scales, S. J., Pepperkok, R., & Kreis, T. E. (1997). Visualization of ER-to-golgi transport in living cells reveals a sequential mode of action for Copii and COPI. Cell, 90(6), 1137–1148. https://doi.org/10.1016/s0092-8674(00)80379-7.

Shomron O, Nevo-Yassaf I, Aviad T, Yaffe Y, Zahavi EE, Dukhovny A, Perlson E, Brodsky I, Yeheskel A, Pasmanik-Chor M, Mironov A, Beznoussenko GV, Mironov AA, Sklan EH, Patterson GH, Yonemura Y, Sannai M, Kaether C, Hirschberg K. COPII collar defines the boundary between ER and ER exit site and does not coat cargo containers. J Cell Biol. 2021 Jun 7;220(6):e201907224. doi: 10.1083/jcb.201907224.

Steven Xijin Ge and others, ShinyGO: a graphical gene-set enrichment tool for animals and plants, *Bioinformatics*, Volume 36, Issue 8, April 2020, Pages 2628–2629, https://doi.org/10.1093/bioinformatics/btz931

Südhof TC, Rothman JE. Membrane fusion: grappling with SNARE and SM proteins. Science. 2009 Jan 23;323(5913):474-7. doi: 10.1126/science.1161748.

Sztul E, Chen PW, Casanova JE, Cherfils J, Dacks JB, Lambright DG, Lee FS, Randazzo PA, Santy LC, Schürmann A, Wilhelmi I, Yohe ME, Kahn RA. ARF GTPases and their GEFs and GAPs: concepts and challenges. Mol Biol Cell. 2019 May 15;30(11):1249–1271. doi: 10.1091/mbc.E18-12-0820.

Takeshita N, Manck R, Grün N, de Vega SH, Fischer R. Interdependence of the actin and the microtubule cytoskeleton during fungal growth. Curr Opin Microbiol. 2014 Aug;20:34–41. doi: 10.1016/j.mib.2014.04.005

Thomas, P. D., Ebert, D., Muruganujan, A., Mushayahama, T., Albou, L., & Mi, H. (2021). PANTHER: Making genome-scale phylogenetics accessible to all. Protein Science, 31(1), 8–22. https://doi.org/10.1002/pro.4218

Tyanova, S., Temu, T., Sinitcyn, P., Carlson, A., Hein, M.Y., Geiger, T., Mann, M., and Cox, J. (2016). The Perseus computational platform for comprehensive analysis of proteomics data. Nat. Methods 13, 731–740. 10.1038/nmeth.3901.

Weigel AV, Chang CL, Shtengel G, Xu CS, Hoffman DP, Freeman M, Iyer N, Aaron J, Khuon S, Bogovic J, Qiu W, Hess HF, Lippincott-Schwartz J. ER-to-Golgi protein delivery through an interwoven, tubular network extending from ER. Cell. 2021 Apr 29;184(9):2412–2429.e16. doi: 10.1016/j.cell.2021.03.035.

Whitney JA, Gomez M, Sheff D, Kreis TE, Mellman I. Cytoplasmic coat proteins involved in endosome function. Cell. 1995 Dec 1;83(5):703–13. doi: 10.1016/0092-8674(95)90183-3.

Wu B, Guo W. The Exocyst at a Glance. J Cell Sci. 2015 Aug 15;128(16):2957–64. doi: 10.1242/jcs.156398.

Yang R, Meyer AS, Droujinine IA, Udeshi ND, Hu Y, Guo J, McMahon JA, Carey DK, Xu C, Fang Q, Sha J, Qin S, Rocco D, Wohlschlegel J, Ting AY, Carr SA, Perrimon N, McMahon AP. A genetic model for in vivo proximity labeling of the mammalian secretome. Open Biol. 2022 Aug;12(8):220149. doi: 10.1098/rsob.220149.

Zhang, C.-J., Bowzard, J.B., Anido, A. and Kahn, R.A. (2003), Four ARF GAPs in *Saccharomyces cerevisiae* have both overlapping and distinct functions. Yeast, 20: 315–330. https://doi.org/10.1002/yea.966

Zhang, Y., Song, G., Lal, N. K., Nagalakshmi, U., Li, Y., Zheng, W., Huang, P., Branon, T. C., Ting, A. Y., Walley, J. W., & Dinesh-Kumar, S. P. (2019). Turboid-based proximity labeling reveals that UBR7 is a regulator of n NLR immune receptor-mediated immunity. Nature Communications, 10(1). https://doi.org/10.1038/s41467-019-11202-z.

